# High-throughput synthesis and specificity characterization of natively paired antibodies using oPool^+^ display

**DOI:** 10.1101/2024.08.30.610421

**Authors:** Wenhao O. Ouyang, Huibin Lv, Wenkan Liu, Ruipeng Lei, Zongjun Mou, Tossapol Pholcharee, Logan Talmage, Meixuan Tong, Yiquan Wang, Katrine E. Dailey, Akshita B. Gopal, Danbi Choi, Madison R. Ardagh, Lucia A. Rodriguez, Xinghong Dai, Nicholas C. Wu

**Author notes:** To whom correspondence should be addressed. (N.C.W.).

## Abstract

Antibody discovery is crucial for developing therapeutics and vaccines as well as understanding adaptive immunity. However, the lack of approaches to synthesize antibodies with defined sequences in a high-throughput manner represents a major bottleneck in antibody discovery. Here, we presented oPool^+^ display, a high-throughput cell-free platform that combined oligo pool synthesis and mRNA display to rapidly construct and characterize many natively paired antibodies in parallel. As a proof-of-concept, we applied oPool^+^ display to probe the binding specificity of >300 uncommon influenza hemagglutinin (HA) antibodies against 9 HA variants through 16 different screens. Over 5,000 binding tests were performed in 3-5 days with further scaling potential. Follow-up structural analysis of two HA stem antibodies revealed the previously unknown versatility of IGHD3-3 gene segment in recognizing the HA stem. Overall, this study established an experimental platform that not only accelerate antibody characterization, but also enable unbiased discovery of antibody molecular signatures.

## INTRODUCTION

Antibodies are central to the immune system for protection against pathogen infection. Therefore, identification of antibodies that target pathogens of interest is key to the understanding of adaptive immunity as well as the development of effective therapeutics and vaccines. In recent years, advances in single-cell B-cell receptor sequencing (scBCR-seq) have greatly improved the capacity to discover novel antibodies^1^. Thousands of natively paired antibody sequences can be obtained from a single scBCR-seq experiment. By contrast, downstream characterization of these antibody sequences remains costly, labor intensive, and time consuming, involving cloning, expression, purification, and testing the binding activities of different antibodies individually. At the same time, protein display technologies offer a high-throughput solution for characterizing antibody binding activity^2^, with antibody library construction being an essential first step. Methods for constructing antibody libraries with random heavy-light chain pairing from B cell repertoires are well-established^3,4^. However, there is a lack of approaches to synthesize custom-made antibody libraries with precise heavy-light chain pairing from a defined list of antibody sequences. This technical barrier has restricted the application of protein display technologies in antibody research, including large-scale characterization of previously discovered antibodies.

Hemagglutinin (HA) is the major antigen of influenza A and B viruses. Influenza A HA is further divided into two groups with a total of 19 antigenic subtypes (H1-H19). Group 1 HA includes H1, H2, H5, H6, H8, H9, H11, H12, H13, H16, H17, H18, and H19, whereas group 2 HA includes H3, H4, H7, H10, H14, and H15. HA is a homotrimeric glycoprotein consisting of a hypervariable globular head domain atop a conserved stem domain^5,6^. The functions of HA are critical for viral entry. The globular head domain engages the sialylated receptor, whereas the stem domain possesses the membrane fusion machinery. In the past two decades, many human antibodies to the HA stem have been discovered and characterized^7–11^. In contrast to HA head antibodies, which are usually strain-specific, HA stem antibodies often cross-neutralize multiple influenza subtypes. Several recurring sequence features have been observed among HA antibodies isolated from different individuals, such as IGHV1-69, IGHV1-18, IGHD6-1, IGHD3-9 for stem antibodies, and IGHV2-70 and IGHD4-17 for head antibodies^7,11–15^. However, many HA antibodies do not contain any known recurring sequence features, suggesting there are additional features yet to be discovered. Discovering the recurring sequence features of HA antibodies is critical for the molecular understanding of antibody responses at the population level, which will in turn benefit the development of universal influenza vaccines.

In this study, we developed oPool^+^ display, a rapid and cost-effective cell-free platform that combined oligo pool synthesis with mRNA display to assemble and screen natively paired antibodies in a highly parallel manner. As a proof-of-concept, we first synthesized a library of 325 natively paired HA antibodies with uncommon germline gene usages, then characterized their binding activity to seven different HAs from influenza A and B viruses as well as an H1 stem construct and an H3 stem construct. We also carried out competition screens against a known HA stem antibody for binding to the same set of HAs, allowing epitope inference. Our screening showed that 114 of the 325 antibodies bound to at least one of the nine HA screened, 45 of which were further identified as HA stem antibody candidates. Extensive experimental validations confirmed the robust performance oPool^+^ display. We then demonstrated oPool^+^ display’s potential in discovering antibody sequence features through structural and functional characterizations of 16.ND.92 and AG11-2F01, two group 1 stem antibody enriched in the screen. 16.ND.92 and AG11-2F01 exhibited distinct binding modes of IGHD3-3 to HA stem, yet both antibodies were broadly reactive and protected against lethal influenza challenge *in vivo*. This observation substantiated that IGHD3-3 is a recurring sequence feature of HA stem antibodies and demonstrated its versatility in binding to HA stem.

## RESULTS

### High throughput assembly of natively paired antibody library via oligo pools

We previously curated a dataset containing 5,561 human monoclonal HA antibodies, 1,082 (19.5%) of which are known to bind to either the head domain or the stem domain^16^. Of the remaining 4,479 (80.5%) HA antibodies which lack epitope information, 292 have complete heavy chain variable (V_H_) and light chain variable (V_L_) sequences available and are not encoded by well-characterized sequence features of HA stem antibodies, namely IGHV1-69, IGHV1-18, IGHV6-1, and IGHD3-9^7,12–14^ (**Figure S1A and Table S1**). These 292 HA antibodies were included in the synthesis of our natively paired antibody library. In addition, three known stem antibodies, namely 31.a.55, AG11-2F01, and 042-100809-2F04^7,17^, as well as 30 known HA head antibodies^6,15,18–25^ were included as controls, bringing the total library size to 325 antibodies **(Figure 1A)**.

**Figure 1.**
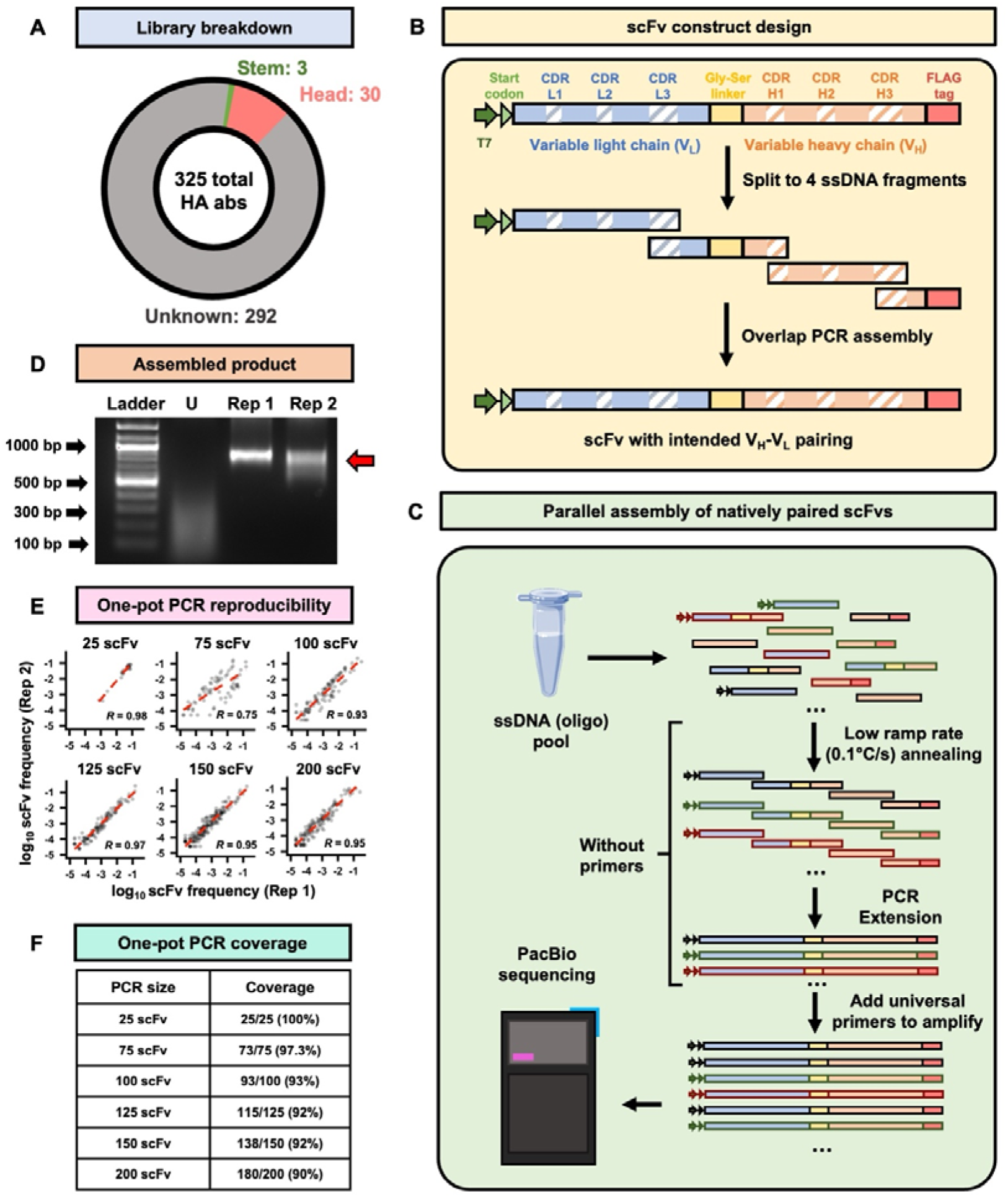
Curation and synthesis of the natively paired HA antibody library. **(A)** The overall breakdown of the HA antibody library. **(B)** Design of oligos for scFv assembly. Each given scFv construct contains a T7 promoter and a start codon at the N-terminal as well as a FLAG tag at the C-terminal. The scFv sequences were then split into 4 fragments at the selected CDR regions, with overlap between adjacent fragments. Through an overlap PCR, oligos of the same construct would preferably anneal to each other, ensuring the assembly of natively paired scFvs. **(C)** Synthesis of the natively paired HA antibody library. Synthesized oligo pools containing scFv fragments were assembled via a two-stage PCR (**see Materials and Methods**). **(D)** The unassembled and assembled oligo pools were compared by agarose gel electrophoresis. “U”: unassembled oligo pool. “Rep1”: replicate 1. “Rep2”: replicate 2. The red arrow indicates the target size (800-900 bp) for full length scFvs. **(E-F)** The reproducibility **(E)** and coverage **(F)** of the scFv assembly using varying numbers of scFv per PCR, ranging from 25 to 200. The Pearson correlation coefficient (*R*) of the occurrence frequencies of individual scFvs between the two replicates were indicated. Micro-tube icon by Servier https://smart.servier.com/ is licensed under CC-BY 3.0 Unported, available via Bio Icons.

To synthesize the natively paired antibody library in a high-throughput manner, we aimed to leverage recent advances in oligo pool synthesis. The maximum length of each oligo in commercial oligo pool synthesis is around 300 to 350 nucleotides. By contrast, the length of a single-chain variable fragment (scFv), which is the smallest format of a human antibody, is around 800 to 900 nucleotides. As a result, each given scFv sequence was split into 4 oligos with overlaps at the diverse complementary-determining regions (CDRs), namely the CDRs H1, H3, and L3 (**Figure 1B-C and Table S2**). We then performed codon randomization to ensure that the overlaps among oligos for the same scFv are unique at the nucleic acid level. This would help prevent mis-annealing between oligos from different scFvs, especially if they shared similar amino acid sequences (**Figure 1B, Material and Methods**). Through an one-pot overlap PCR, full-length scFv sequences could then be generated with the intended native V_H_ and V_L_ pairing. Subsequently, this strategy was applied to design the oligo pools for our library of 325 antibodies (**Figure 1C-F, Figure S1B-C, and Table S2**).

The length of the assembled product was consistent with that of full-length scFvs (**Figure 1D**). To thoroughly evaluate the effectiveness of our synthesis strategy, we then performed several assembly PCRs with the starting oligos of 25, 75, 100, 125, 150, and 200 scFvs each in a single tube (**Figure 1E-F and Figure S2**). PacBio sequencing revealed the high reproducibility of the one-pot PCR assembly across the board, with a Pearson correlation coefficient of 0.95 even at 200 scFvs per PCR (**Figure 1E**). The one-pot PCR assembly strategy also achieved high coverage of the targeted antibodies, with only a subtle drop from 100% to 90% as the complexity of the PCR increased from 25 to 200 scFvs (**Figure 1F**). For our final library of all 325 scFvs, a Pearson correlation coefficient of 0.83 was observed between replicates, with a coverage of 322 of 325 (99.1%) natively paired antibodies (**Figure S3**). While we opted to assemble this antibody library through 13 PCRs with 25 scFvs each tube, our results demonstrated the potential to further increase the throughput of our antibody library synthesis strategy by at least one orders of magnitude.

### Rapid specificity characterization of the natively paired antibodies by mRNA display

To characterize antibody specificity from our natively paired scFv library, we utilized mRNA display^26–28^, a well-established cell-free screening approach that allows rapid screening for protein binders. Briefly, each scFv was covalently linked to the RNA molecule that encoded it, thus providing a phenotype-genotype linkage (**Figure S4A, Material and Methods**). The mRNA-displayed antibody library was then selected against seven HAs from different influenza A subtypes and influenza B strains, namely H1N1 A/Solomon Island/3/06 (H1/SI06), H1N1 A/Michigan/45/2015 (H1/MI15), H3N2 A/Singapore/INFIMH-16-0019/2016 (H3/SP16), H5N1 A/ Qinghai/1A/2005 (H5/QH05), H7N9 A/Shanghai/2/2013 (H7/SH13), B/Phuket/3073/2013 (B/Phu13), and B/Lee/1940 (B/Lee40). Selections were also performed against two HA stem domain constructs that were designed based on H1N1 A/Brisbane/59/2007 HA and H3N2 A/Finland/486/2004 HA, respectively^29,30^ (**Figure 2 and Figure S4**). After one round of selection, the pre- and post-selection libraries were then analyzed by PacBio sequencing to quantify the enrichment of each scFv (**Table S3**). Pearson correlation coefficients ranging from 0.58 to 0.86 were observed between biological replicates (**Figure S5**), demonstrating the reproducibility of the selections. In total, 114 of the 325 scFvs were identified to target one of the HAs screened, with 52 targeting more than one HA and 11 targeting more than two HAs (**Table S4**). All stem antibody controls were enriched in at least one screens, while only 17 of the 30 head antibody controls were enriched, likely due to the considerably limited binding breadth of head antibodies. Moreover, the stem antibody controls were highly enriched in the selections against the HA stem domain constructs, whereas the head antibody controls were not, further validating that selection took place.

**Figure 2.**
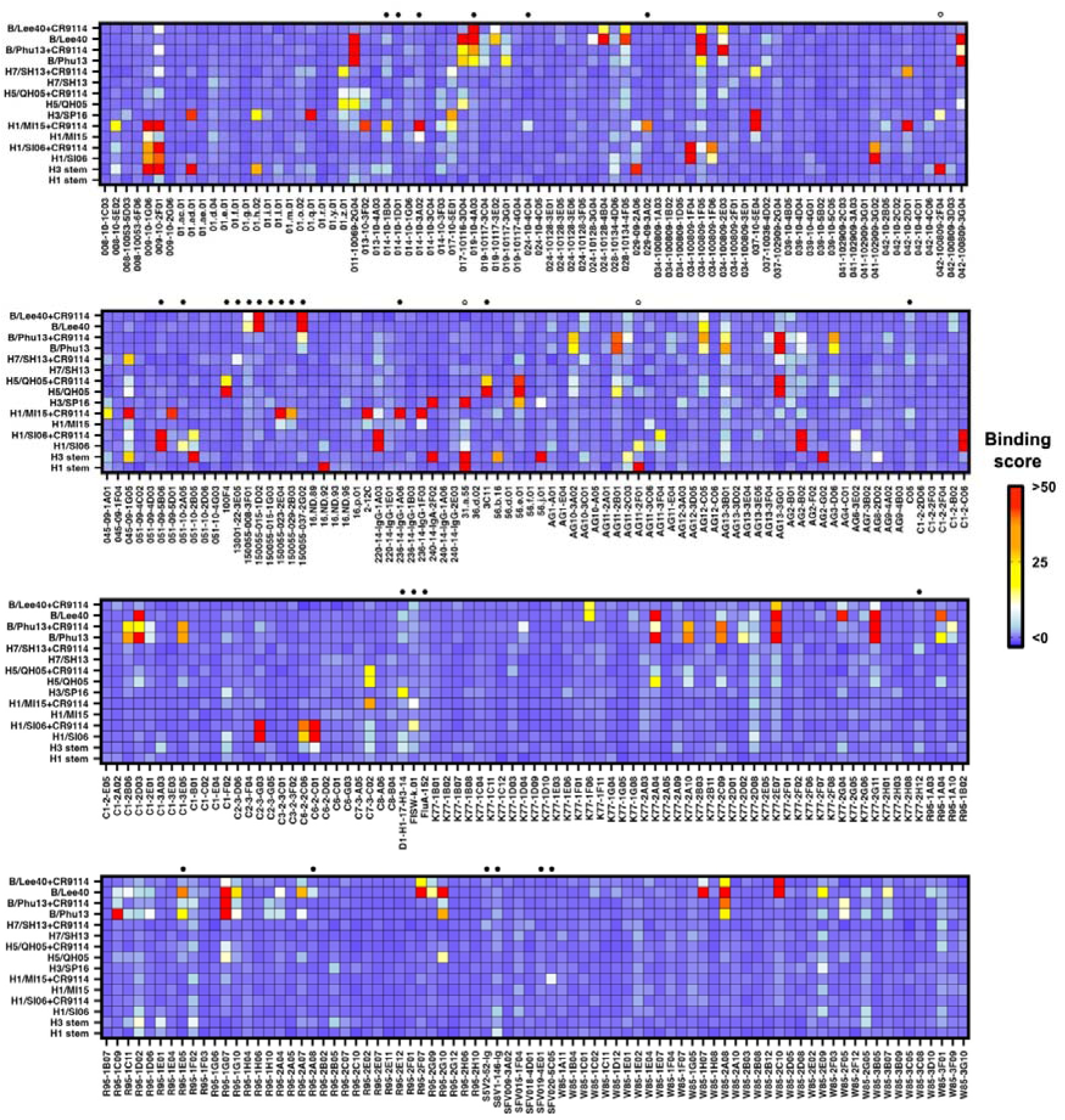
Rapid specificity characterization of the natively paired antibodies by mRNA display. The screening results of each scFv are shown as a heatmap. *X*-axis represents the scFv names. *Y*-axis represents each individual screens. Screens against CR9114 IgG pre-bound HAs are indicated by “+CR9114”. Binding scores shown were adjusted with robust scaling. Individual cutoffs were set for each screen to determine positive hits (**Table S4, Materials and Methods**). Black circles at the top of the heatmap indicate head antibody controls, whereas empty circles indicate stem antibody controls.

We also performed seven competition screens against CR9114^10^, a broadly neutralizing HA stem antibody, for binding to H1/SI06, H1/MI15, H3/SP16, H5/QH05, H7/SH13, and B/Phu13, and B/Lee40 (**Figure 2 and Figure S6**). Competition screens were performed using the same protocol as the mRNA display selections described above, except that CR9114 was pre-bound to the HAs. While the CR9114 competition screen for H3/SP16 did not yield consistent PCR bands post-selection, all six other screens resulted in reproducible bands and were analyzed by PacBio sequencing. Of these six screens, five had Pearson correlation coefficients of >0.7 between replicates (**Figure S7**), demonstrating their reproducibility. The results of our competition screening were in agreement with previous studies. For example, two of our stem antibody controls, AG11-2F01 and 31.a.55, competed with CR9114 in our screens, which is consistent with previous studies of these two antibodies^7,17^. Additionally, 15 of the 17 (88%) HA head antibody controls targeting at least one HA in our screens did not compete with CR9114, further supporting the effectiveness of the competition screen (**Table S4**). Together, we identified five antibody candidate targeting group 1 HA stem, 12 targeting group 2 HA stem, as well as 28 targeting influenza B HA stem (**Figure S5 and Table S4**). Of note, one of the H3 stem antibody candidates, AG2-G02, was concurrently identified by a machine learning approach and experimentally confirmed in another study of ours^16^.

### oPool^+^ display demonstrates robust performance in specificity characterization

To systematically evaluate the performance of oPool^+^ display, we used both biolayer interferometry (BLI) and ELISA to validate the binding activities of selected antibodies (**Figure 3**). In brief, 25 antibodies were selected, recombinantly expressed and purified in fragment antigen-binding (Fab) and IgG formats. Their binding activities were then tested against all nine HAs used in our screens (**Figure 3A-B, Figure S8**). This validation experiment substantiated the robust performance of oPool^+^ display. A true positive rate of 80.4% (41/51) and a true negative rate of 95.4% (166/174) was observed in BLI validation with Fabs, while a true positive rate of 71.7% (43/60) and a true negative rate of 96.4% (159/165) was observed in ELISA validation with IgG (**Figure 3C**). Of note, both experimental validations indicated that the majority of the false positive and false negative results (11/18 for BLI, 14/23 for ELISA) were from three screens (H3 stem, H1/SI06, and H1/MI15). To examine the influences of different antibody formats on binding, we determined the dissociation constant (K_D_) of three H1 stem antibodies and six H3 stem antibodies that were enriched in our screen in both Fab and scFv formats (**Figure 3D, Figure S9 and S10**). All nine antibodies retained binding in both formats, with no apparent correlation between the K_D_ values. However, some of the false negatives could still be due to weakened binding of the antibody in scFv format during the screens.

**Figure 3.**
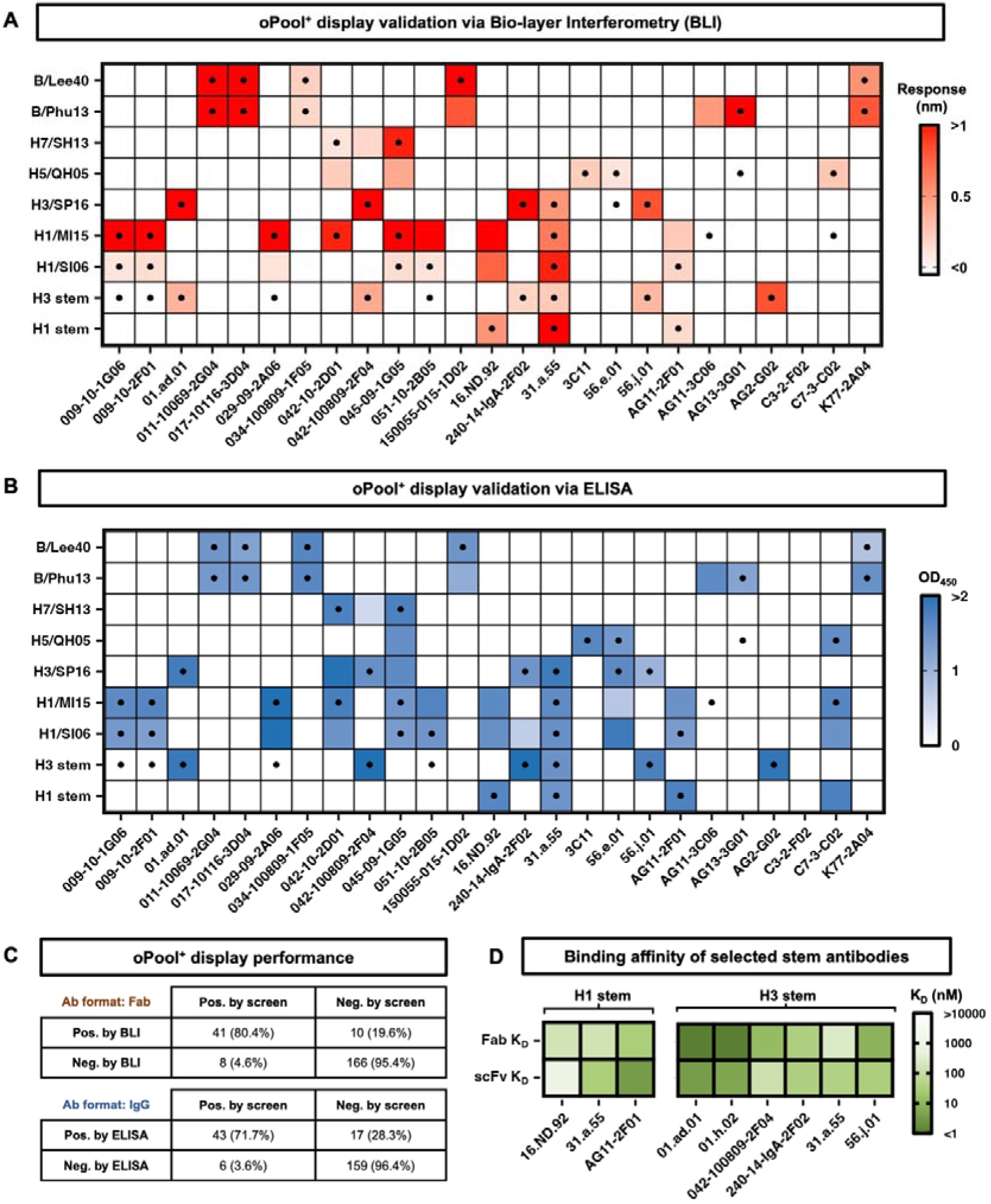
Systematic validation of oPool^+^ display. (A-B) Binding activity of 25 antibodies against different HAs was measured by **(A)** biolayer interferometry (BLI) with antibodies in Fab format and **(B)** ELISA with antibodies in IgG format. **(A)** The response signals during the association phase and **(B)** the OD_450_ values were shown as heatmaps. The dots represent the hits in oPool^+^ display. X-axis represents antibody names. Y-axis represents antigen names. **(C)** Binary confusion matrices based on BLI and ELISA validations are shown. **(D)** Binding affinity of selected HA stem antibody candidates in both scFv and Fab format. Their dissociation constants (K_D_) against H1 stem and H3 stem are shown as heatmaps. Of note, 31.a.55 was a positive control for binding to both H1 stem and H3 stem^7^, AG11-2F01 was a positive control for binding to H1 stem^17^, and 042-100809-2F04 was a positive control for binding to H3 stem^54^.

We then further assessed the performance of our CR9114 competition screens using 16 of the 25 antibodies validated above (**Figure 4A-B**). To quantify the competition, we first defined the CR9114 competition index as the log fold change of enrichments with versus without the presence of CR9114 during screen (**Table S4, Figure S11, Materials and Methods**). A higher CR9114 competition index would indicate overlapping epitopes between the scFv and CR9114, while a lower CR9114 competition index would indicate opposite (**Figure 4A**). Experimental validation of these competition indices was then performed using BLI, where the ratio of binding responses to HA with and without CR9114 pre-bound was quantified (**Materials and Methods**). Pearson correlation of 0.66 was observed between the competition indices and the validation results (**Figure 4B, Figure S12, and Table S5**), confirming the reliability of our CR9114 competition screens. Together, our results showed that oPool^+^ display enabled rapid and accurate specificity characterization of natively paired antibodies.

**Figure 4.**
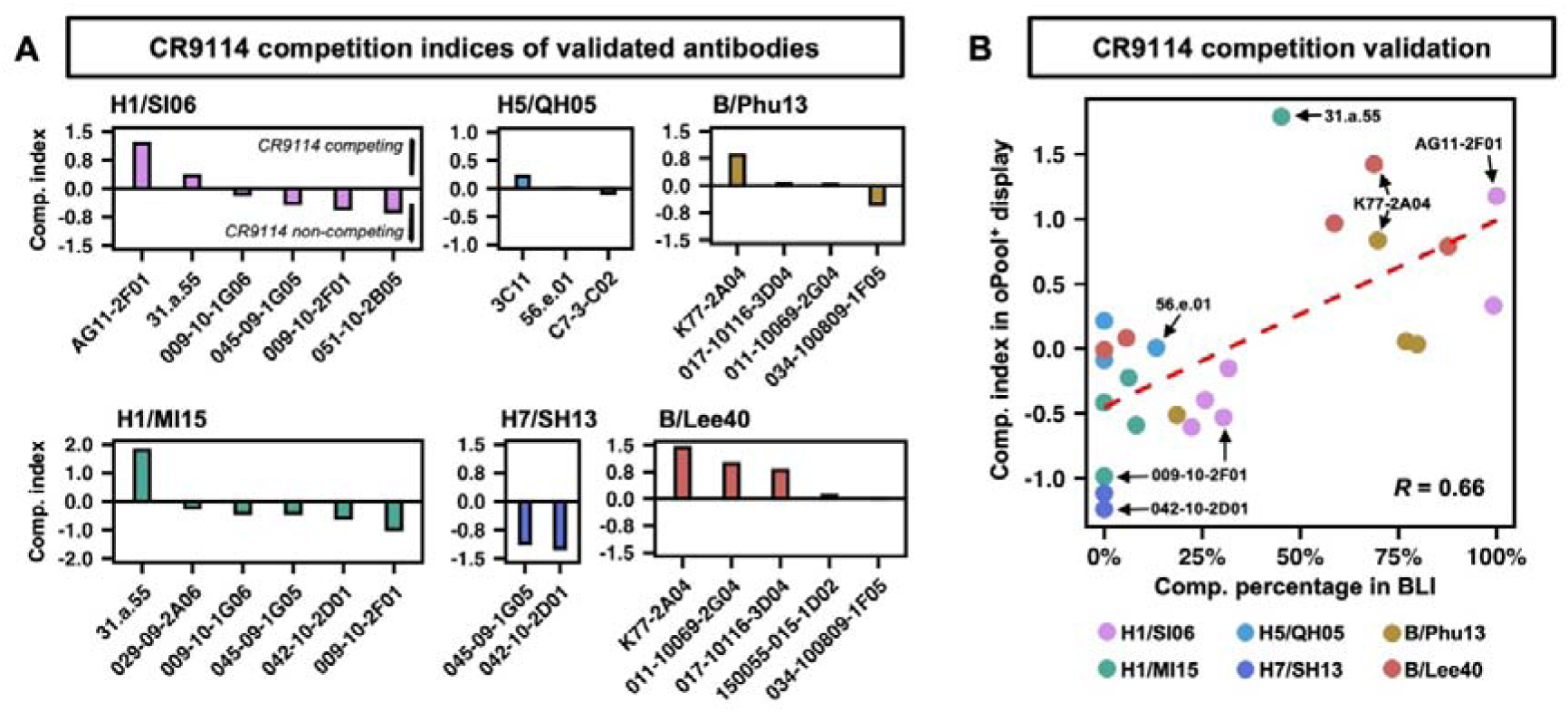
CR9114 competition profile of validated antibodies. **(A)** CR9114 competition indices of validated antibodies. The competition indices calculated from oPool^+^ display are shown. High positive values indicate CR9114 competition, low positive and high negative values indicate no CR9114 competition. **(B)** CR9114 competition of validated antibodies based on BLI results, shown as scatter plot against the competition indices. The response of each antibody binding to the antigen were measured during the association step with or without prior saturation binding of CR9114. The percentage of competition is shown with the range of 0% (no competition) to 100% (complete competition). Pearson correlation coefficient (*R*) between replicates is indicated. The linear fit lines are shown as red dash lines. Representative antibodies are labelled.

### AG11-2F01 and 16.ND.92 have similar sequence features but distinct binding modes

One of the H1 stem antibodies identified from our screen was 16.ND.92, which was originally isolated from a young individual in an H5N1 influenza vaccine trial^7^. 16.ND.92 was encoded by IGHV3-74/IGHD3-3/IGKV1-5 (**Table S4**). Coincidentally, both IGHD3-3 and IGKV1-5 were utilized by AG11-2F01, which was one of the two positive controls against H1 stem in our screen (**Figure 2 and Figure S5**). Moreover, 16.ND.92 and AG11-2F01 shared a similar FG[V/L] motif encoded by the reading frame +3 of IGHD3-3 (**Figure S13**). This observation led us to hypothesize that 16.ND.92 and AG11-2F01 engaged the HA stem via similar binding modes. Consequently, we determined the cryo-EM structures of H1N1 A/Solomon Islands/03/2006 (H1/SI06) HA in complex with AG11-2F01 and 16.ND.92 to resolutions of 2.89 Å and 2.82 Å, respectively (**Figure 5A-E and Table S6**). Contrary to our hypothesis, the cryo-EM structures revealed very different binding modes between the two antibodies. While AG11-2F01 bound to HA stem horizontally, 16.ND.92 had a downward approaching angle towards HA (**Figure 5A**). Relatedly, the epitope of 16.ND.92 shifted slight upward compared to that of AG11-2F01 (**Figure 5B**).

**Figure 5.**
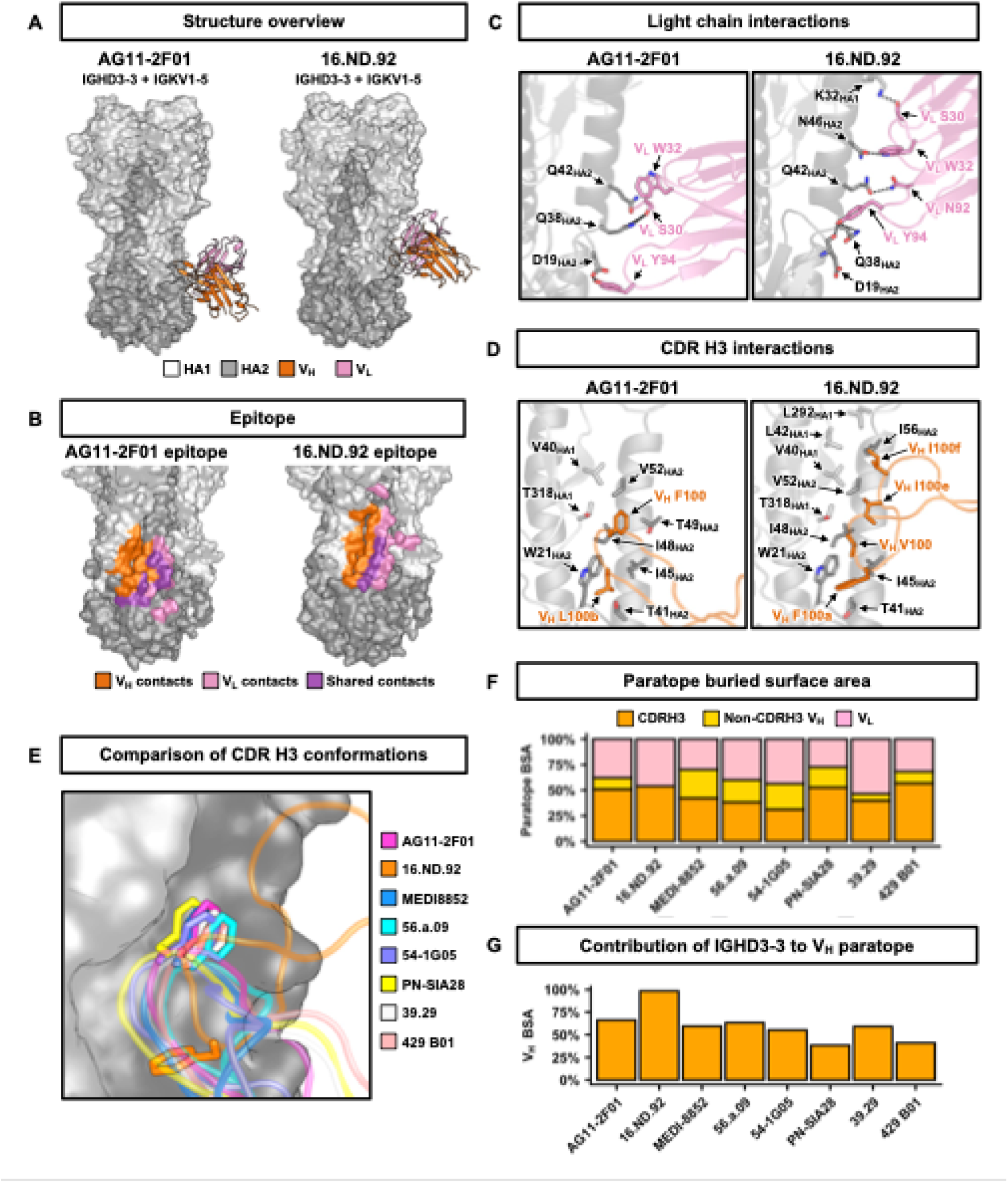
Structural analysis of AG11-2F01 and 16.ND.92. **(A)** Cryo-EM structures of AG11-2F01 and 16.ND.92 in complex with H1/SI06 HA. HA1 is in light grey. HA2 is in dark grey. Heavy chain variable domain (V_H_) is in orange. Light chain variable domain (V_L_) is in pink. **(B)** Epitopes of AG11-2F01 and 16.ND.92. V_H_ contacts are in orange. V_L_ contacts are in pink. Contacts shared by both V_H_ and V_L_ are in purple. **(C)** Interactions between light chain and HA are shown. H-bonds are represented by black dashed lines. **(D)** Interactions between CDR H3 and HA are shown. **(E)** Overlay of the CDR H3 loops from IGHD3-3 HA-stem antibodies. HA is in surface representation. **(F)** Contributions of CDR H3 (orange), non-CDR H3 V_H_ (yellow), and V_L_ (pink) to the paratope buried surface areas (BSA) of the indicated antibodies. **(G)** Contributions of IGHD3-3-encoded residues to V_H_ paratope BSA of the indicated antibodies.

Although both 16.ND.92 and AG11-2F01 were encoded by IGKV1-5, their light chains interacted with the HA stem differently. For example, V_L_ S30 of AG11-2F01 H-bonded with HA2 Q38, whereas that of 16.ND.92 H-bonded with HA1 K32 (**Figure 5C**). Similarly, despite sharing an FG[V/L] motif in their IGHD3-3-encoded regions, 16.ND.92 and AG11-2F01 used this motif to interact with different parts of the HA stem (**Figure 5D**). For the FGL motif in AG11-2F01, V_H_ F100 inserted into a hydrophobic pocket in the HA stem centering at HA2 I48, whereas V_H_ L100b inserted into a lower pocket centering at HA2 W21. By contrast, this lower pocket was occupied by the V_H_ F100a of the FGV motif in 16.ND.92. The paratope of 16.ND.92 also involved IGHD3-3-encoded V_H_ V100, I100e, and I100f, allowing its CDR H3 to bind to upper pockets in the HA stem that were not engaged by AG11-2F01 (**Figure 5D**). Together, our structural analyses showed that AG11-2F01 and 16.ND.92 formed distinct molecular interactions with HA stem.

### 16.ND.92 utilizes IGHD3-3 in a unique manner for binding to HA stem

Previous studies have determined the structures of several HA stem antibodies with an FG[V/L/I] motif in the CDR H3 that was encoded by reading frame +3 of IGHD3-3, including MEDI8852, 56.a.09, 54-1G05, 39.29, PN-SIA28, and 429 B01^7,9,31–34^. These six HA stem antibodies used either IGHV6-1 or IGHV3-30, unlike AG11-2F01 and 16.ND.92, which used IGHV4-38-2 and IGHV3-74, respectively. Nevertheless, the CDR H3 conformation of AG11-2F01 resembled that of MEDI8852, 56.a.09, 54-1G05, 39.29, PN-SIA28, and 429 B01 (**Figure 5E**). Moreover, the IGHD3-3-encoded FG[V/L/I] motifs of these seven antibodies bound to the HA stem in a similar fashion (**Figure 5E**). In comparison, the CDR H3 conformation of 16.ND.92 was different due to more extensive involvement of IGHD3-3 in binding (**Figure 5D-E**).

The unique usage of IGHD3-3 for binding enables 16.ND.92 V_H_ to interact with the HA stem exclusively through CDR H3, whereas the V_H_ paratopes of other IGHD3-3 HA stem antibodies involved non-CDR H3 regions (**Figure 5F and Table S7**). Similarly, IGHD3-3 accounted for 98.6% of the buried surface area of the 16.ND.92 V_H_ paratope, but 38% to 63% of the V_H_ paratopes of other IGHD3-3 HA stem antibodies (**Figure 5G and Table S7**). These observations not only substantiated that reading frame +3 of IGHD3-3 was a recurring sequence feature of HA stem antibodies, but also demonstrated that it could pair with diverse IGHV genes and interact with HA stem via different binding modes.

### AG11-2F01 and 16.ND.92 are neutralizing antibodies with *in vivo*protection activity

Given the different binding modes of AG11-2F01 and 16.ND.92, we further aimed to compare their binding breath, *in vitro* neutralization, and *in vivo* protection activity. ELISA showed that both AG11-2F01 and 16.ND.92 bound to all H1 and H5 HAs tested (**Figure 6A and Figure S14)**, Microneutralization assay against six H1N1 strains further revealed their neutralizing activity (**Figure 6B**). Both AG11-2F01 and 16.ND.92 also protected mice against a lethal challenge of H1N1 A/Puerto Rico/8/1934 (PR8), based on the weight loss profiles (**Figure 6C-D**), survival analyses (**Figure 6E-F**), and lung viral titers at day 3 post-infection (**Figure 6G-H**). Nonetheless, our results indicated that the *in vivo* therapeutic protection activity of 16.ND.92 was stronger than AG11-2F01. While only 20% (1/5) of the mice therapeutically treated with AG11-2F01 survived (**Figure 6E**), 80% (4/5) of the mice therapeutically treated with 16.ND.92 survived (**Figure 6F**). Additionally, at day 3 post-infection, lung viral titers of mice therapeutically treated with 16.ND.92 were ∼15-fold lower than those treated with AG11-2F01 (**Figure 6G-H**). Notably, 16.ND.92 had comparable, if not weaker, *in vitro* neutralizing activity than AG11-2F01 against PR8 (**Figure 6B**). Consequently, the stronger *in vivo* protection activity of 16.ND.92 against PR8 may at least be partly attributed to its more downward approaching angle to the HA stem (**Figure 5A**), which could help position the Fc region closer to effector cells.

**Figure 6.**
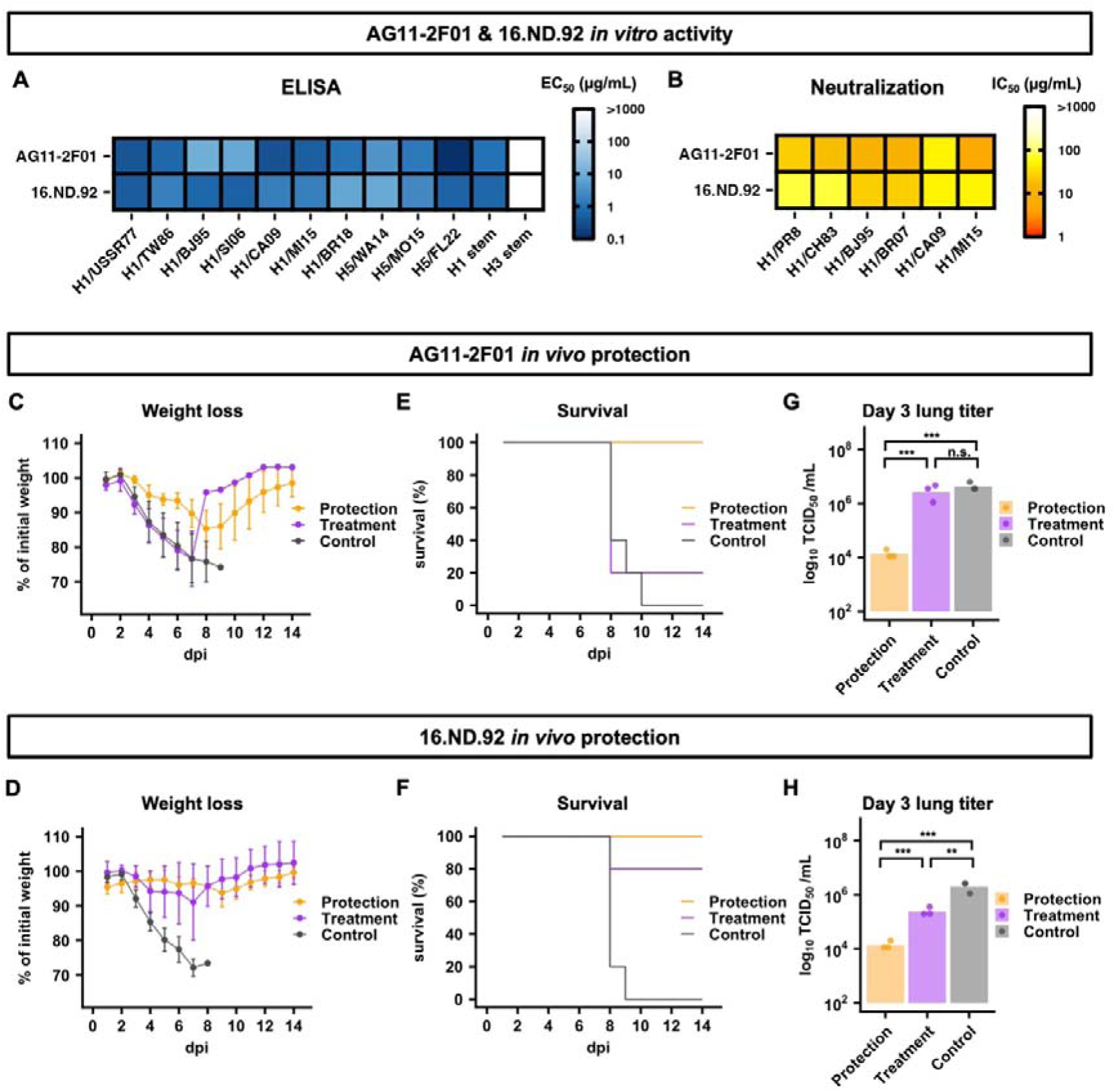
*in vitro* and *in vivo* protection of AG11-2F01 and 16.ND.92. **(A)** The binding activities of AG11-2F01 and 16.ND.92 against recombinant HA proteins from the indicated H1 and H5 strains were measured by ELISA. The EC_50_ values are shown as a heatmap. **(B)** The neutralization activity of AG11-2F01 and 16.ND.92 against different recombinant H1N1 viruses was measured by a microneutralization assay. The IC_50_ values are shown as a heatmap. **(A-B)** Strain names are abbreviated as follows: H1N1 A/Puerto Rico/8/1934 (H1/PR8), H1N1 A/USSR/90/1977 (H1/USSR77), H1N1 A/Chile/1/1983 (H1/CH83), H1N1 A/Taiwan/01/1986 (H1/TW86), H1N1 A/Beijing/262/1995 (H1/BJ95), H1N1 A/Solomon Island/3/2006 (H1/SI06), H1N1 A/Brisbane/59/2007 (H1/BR07), H1N1 A/California/04/2009 (H1/CA09), H1N1 A/Michigan/45/2015 (H1/MI15), H1N1 A/Brisbane/02/2018 (H1/BR18), H5N8 A/northern pintail/WA/40964/2014 (H5/WA14), H5N2 A/snow goose/Missouri/CC15-84A/2015 (H5/MO15), and H5N1 A/bald eagle/Florida/W22-134-OP/2022 (H5/FL22). H1 stem and H3 stem represents the HA stem constructs designed based on H1N1 A/Brisbane/59/2007 HA and H3N2 A/Finland/486/2004 HA, respectively^29,30^. **(C-H)** The *in vivo* protection activity of AG11-2F01 and 16.ND.92 against lethal challenge of PR8 virus was assessed by **(C-D)** weight loss profiles, **(E-F)** Kaplan-Meier survival curves, and **(G-H)** lung viral titers at day 3 post-infection. *P* values were computed by two-tailed student’s t-test. ***: *P* < 0.001; **: *P* < 0.01; n.s.: not significant.

## DISCUSSION

Antibody discovery has led to significant advancements on many fronts, including antibody-based therapeutics as well as vaccine designs^11,35–38^. Discovery of natively paired antibody sequences has been hugely accelerated by scBCR-seq in the past few years^1^. However, going from sequence information to specificity characterization remains a major bottleneck in antibody discovery. In this study, we presented oPool^+^ display, an experimental platform that allows specificity characterization of antibodies with defined sequences in a highly parallel fashion. Importantly, oPool^+^ display was more cost-efficient (∼$30 per antibody) and faster (∼3-5 days) than the conventional methods that require cloning and recombinant expression of individual antibodies (∼$200-350 per antibody, weeks to months) (**Table S8**). As a proof-of-concept, we applied oPool^+^ display to delineate the binding specificity of hundreds of HA antibodies that were left uncharacterized in the literature. Follow-up analysis of AG11-2F01 and 16.ND.92 further revealed the versatility of IGHD3-3 in targeting the HA stem.

A key feature of oPool^+^ display is its relatively simple protocol. A previous study has shown that the throughput for screening antibodies with defined sequences can be increased by using liquid handlers to express individual antibodies one by one^39^. In comparison, oPool^+^ display uses a near one-pot strategy for antibody library synthesis and screening. As suggested by our results, one 96-well plate PCR can allow rapid assembly of library up to ∼20,000 natively paired antibodies. With current display technologies, such as mRNA display, screening of the library against 10 to 20 antigens can be done by one person in hours. Moreover, it only requires standard benchtop equipment commonly found in a regular molecular biology lab. In addition, the constructed library can be stored long-term as DNA for future use when new antigens of interest emerge, such as novel HA variants or subtypes. The library can also be expanded by merging with additional oligo pools when new panels of antibodies are discovered. Therefore, oPool^+^ display not only bridges the gap between scBCR-seq and protein display technologies through massively parallel reconstruction of sequenced antibodies, but also provide more flexibility of the current antibody discovery pipelines. After the natively paired antibody sequences are obtained from scBCR-seq, oPool^+^ display can be applied to validate and characterize the specificity of a large panel of antibody candidates at any time. The synergy between oPool^+^ display and scBCR-seq can streamline the transition from antibody discovery to antibody characterization.

Previous studies have shown that IGHV6-1 and IGHV3-30 HA stem antibodies often utilize IGHD3-3-encoded FG[V/L/I] motif for binding to HA stem^7,9,31–34^. As demonstrated by our work here, IGHD3-3-encoded FG[V/L/I] motif can also pair with other IGHV genes to target HA stem, substantiating that it is an IGHV-independent recurring sequence feature of HA stem antibodies. Our results further revealed that IGHD3-3 can engage the HA stem via different binding modes. These observations are comparable to those of IGHD3-9, which is utilized by HA stem antibodies with various IGHV genes and can bind to HA stem in two different reading frames^14^. Similarly, recent studies have identified IGHD3-22 as an IGHV-independent recurring sequence feature of antibodies that target a conserved site on SARS-CoV-2 spike^40,41^. Although antibody sequence analysis typically focuses on IGHV genes, the contribution of IGHD genes to antibody responses should not be overlooked since emerging evidence suggests that it might be more important than previously thought.

Although this study focused on influenza HA stem as a proof-of-concept, oPool^+^ display can be generalized to any antigens of interest as long as they can be recombinantly purified. Importantly, oPool^+^ display can be leveraged for epitope mapping, given that it enables competition screening. Such application will be particularly valuable for antigens that have largely unknown antigenicity but have several antibodies with known epitopes, such as those from emerging pathogens^42,43^. Furthermore, the capability of constructing custom-made antibody libraries means that oPool^+^ display has the potential to benefit the development of machine learning models for antibody engineering, specificity prediction, and *de novo* design, as a major throughput bottleneck still exists in experimental validation^16,44–46^. We envision that prediction results from these models can be rapidly validated by oPool^+^ display, which will in turn facilitate iterative refinement of the models for more advanced applications.

We acknowledge that oPool^+^ display has some limitations. First, oPool^+^ display requires antibodies to be presented as scFvs, which may lose functionality compared to its Fab counterpart^47–49^. Second, antibodies with a fast off-rate may result in false negatives in oPool^+^ display, since it depends on monovalent binding. Inadequate wash during selection could also partially explain the false positive hits in our results. Nonetheless, a few solutions can be adopted in future studies. Performing multiple rounds of mRNA display selection, or replacing it with other protein display technologies that support multivalent binding, such as yeast display and phage display, can further reduce false negative rate^50,51^. In addition, a more stringent wash protocol can potentially reduce false positive rate. As the length of oligo pool synthesis continues to improve, the cost and complexity of oPool^+^ display will further decrease. Overall, we believe that oPool^+^ display represents a starting point for the future advancement of high-throughput approaches to characterize antibodies.

## MATERIALS AND METHODS

### Selection of HA antibodies for paired antibody library synthesis

Members of the natively paired antibody library were selected from a previously curated dataset containing 5,561 human monoclonal antibodies to influenza HA from 60 research publications and three patents^16^. Filters were applied to exclude antibodies that 1) had incomplete sequence information, 2) utilized germline genes that were regarded as recurring sequence features of HA stem antibodies, namely IGHV1-69, IGHV6-1, IGHV1-18 and IGHD3-9^7,12–14^, and 3) were members of known HA stem antibody clonotypes. This resulted in 292 antibody sequences from 7 publications^7,17–19,23,52,53^. Three HA stem antibodies, namely 31.a.55, AG11-2F01, and 042-100809-2F04, as well as 30 HA head antibodies were randomly selected as positive and negative controls^6,7,15,17–25^. Of note, both AG11-2F01 and 042-100809-2F04 were not previously labeled as an HA stem antibody in the curated dataset^16^. Through literature search, AG11-2F01 was found to compete with CR9114^10^, which is an HA stem antibody, for binding to H1^17^, while 042-100809-2F04 was found to bind to group 2 HA stem domain^54^.

### Computational design of the oligo pool sequences for assembly

A summary of the computational design pipeline is described below and summarized in **Figure S1.**

#### Sequence preparation

Selected antibody sequences were first annotated using abYsis^55^. Any missing nucleotides at the 5’ and/or 3’ ends were then filled in using the sequences from the IGHV and IGHJ genes that had the highest identity with the given antibody.

#### Codon randomization and pool assignment of the antibodies

To decrease undesired assembly during antibody library synthesis, codon randomization of each selected antibody sequence was first performed to reduce nucleotide sequence similarity among different antibodies. For a given amino acid, codon usages <15% in *Escherichia coli* were removed from consideration to prevent low translation efficiency during RNA display. For 325 antibodies, total of 2 million randomized sequences were generated to maximize downstream sequence differentiation. All antibody sequences were then split into 8 segments, followed by clustering using CD-HIT^56^. The clusters were generated using the criterion of 70% sequence identity (at least 30% differences between each cluster). The antibody sequences were then reconstructed by selecting necessary segments from different clusters, followed by removing the used clusters. Such reconstruction was repeated until a complete pool (total of 25 antibody sequences) had been reassembled. The deleted clusters were then added back with the reassembled antibody sequences removed. The process was then iterated to generate the remaining pools.

#### Selection of overlap regions and generation of the final oligo pools

Upon the pool assignment of antibodies, the ideal overlap regions were searched over CDR L3, H1, and H3 of each antibody sequence. Six 30-nucleotide long sequences were extracted from each region through frame shifting, then aligned to the complete antibody sequences in the corresponding pool using BLAST+^57^. For each antibody, the overlap sequences that are least similar to other antibodies in the pool were selected. The antibody sequences were then split at the overlap region, followed by the addition of universal adaptor regions at fragments encoding the N-terminal and C-terminal of the antibodies, leading to the generation of the final oligo pools.

### Overlap PCR assembly of the natively paired antibody library

A total of 13 oligo pools were synthesized (Integrated DNA Technologies). The lengths of oligos ranged from around 180 to 330 nucleotides. Each oligo pool contained 100 oligos resuspended in 200 µL water. An assembly PCR was set up for each oligo pool using 1,600 ng of oligos as input. The assembly was performed using KAPA HiFi HotStart ReadyMix (Roche) and a Mastercycler nexus GX2 (Eppendorf). PCR was set up in the absence of primers. From cycles 1-40, PCR was performed with minimal ramp rate (0.1°C/s) in between the denaturing (98°C, 20 s) and annealing steps (62°C, 15 s) to reduce erroneous annealing events. After cycle 40, a universal forward primer 5’-TTC <u>TAA TAC GAC TCA CTA TAG</u> GGA CAA TTA CTA AAG GAG TAT CC-3’ and a universal reverse primer 5’-GGA GCC GCT ACC <u>CTT ATC GTC GTC ATC CTT GTA ATC</u> GGA TCC T-3’ were added to the PCR. The underlined region in the forward primer sequence is the T7 promoter, whereas the underlined region in the reverse primer sequence encodes a FLAG tag. Subsequently, another 15 cycles of PCR were performed to amplify the assembled product. The final PCR product was purified using a Monarch Gel Extraction Kit (New England Biolabs). Two replicates of the assembly were performed.

### Preparation of the biotinylated H1 stem and H3 stem constructs

The H1 stem (mini-HA #4900)^29^, H3 stem (H3ssF)^30^, as well as the seven HA ectodomain constructs were cloned into a customized baculovirus transfer vector. Both constructs contained a N-terminal gp67 signal peptide at the N-terminus as well as a BirA biotinylation site, a thrombin cleavage site, a trimerization domain and a 6xHis-tag at the C-terminus. Recombinant bacmid DNA that carried the H1 stem construct or H3 stem construct was generated using the Bac-to-Bac system (Thermo Fisher Scientific) per manufacturer’s instructions. Baculovirus was generated by transfecting the purified bacmid DNA into adherent Sf9 cells using Cellfectin reagent (Thermo Fisher Scientific) per manufacturer’s instructions. The baculovirus was further amplified by passaging in adherent Sf9 cells at a multiplicity of infection (MOI) of 1.

Recombinant HA constructs were then expressed using 1L of suspension Sf9 cells at an MOI of 1. At day 3 post-infection, Sf9 cells were pelleted by centrifugation at 4,000 xg for 25 min. Soluble recombinant HA constructs were purified from the supernatant by affinity chromatography using Ni Sepharose excel resin (Cytiva) and then size exclusion chromatography using a HiLoad 16/100 Superdex 200 prep grade column (Cytiva) in 20 mM Tris-HCl at pH 8.0, and 100 mM NaCl. The purified protein was concentrated by an Amicon spin filter (Millipore Sigma) and filtered by a 0.22 mm centrifuge Tube Filter (Costar). The purified HA constructs were then biotinylated using a Biotin-Protein Ligase-BIRA kit (Avidity) according to the manufacturer’s instructions. The biotinylated proteins were then purified again through size exclusion chromatography as described above. The A280 absorbance values were measured using a Nanodrop One (Thermo Fisher Scientific) to quantify the protein concentration.

### Antibody screening using mRNA display

The mRNA display was performed based on the protocols provided by previous studies^27,28,58^ with slight modifications.

#### Generation of the puromycin-conjugated mRNA templates

The DNA library was first transcribed by a MEGAscript T7 Transcription Kit (Thermo Fisher Scientific) and purified by a MEGAclear Transcription Clean-Up Kit (Thermo Fisher Scientific) according to manufacturer’s instructions. Ligation was performed using 1 nmol of the mRNA product, 1.1 nmol of the splint oligo (5’-TTT TTT TTT TTT GGA GCC GCT ACC-3’), and 1.2 nmol of the puromycin linker (5’-/5Phos/-d(A)21-(C_9_)3-d(ACC)-puromycin-3’) by the T4 DNA ligase (New England Biolabs) in a 100 µL reaction for 1 hour at room temperature, followed by Lambda exonuclease (New England Biolabs) digestion for 30 mins at 37°C. The puromycin-conjugated mRNA product was purified using a Dynabeads mRNA DIRECT Purification Kit (Thermo Fisher Scientific), aliquoted, and stored at -20°C until used.

#### Preparation of the mRNA-scFv fusion library

The puromycin-conjugated mRNA templates were translated using a PURExpress In Vitro Protein Synthesis Kit (New England Biolabs) with the addition of PURExpress Disulfide Bond Enhancer (New England Biolabs) for 1 hour at 37°C. The reaction was then incubated with 500□mM KCl and 60□mM MgCl_2_ for at least 30 mins at room temperature to promote fusion between the translated scFv and puromycin. EDTA was then added to dissociate ribosomes. The full-length mRNA-scFv product was then purified using Anti-FLAG M2 Magnetic Beads (Millipore Sigma) followed by elution using 3xFLAG peptides (GlpBio). Subsequently, the purified mRNA-scFv product was reverse transcribed using SuperScript IV reverse transcriptase (Thermo Fisher Scientific). The cDNA/mRNA-scFv product was referred as the “pre-selection library”.

#### Preparation of the magnetic beads coated with biotinylated HAs

Biotinylated HA constructs were coated onto the Dynabeads M280-straptavidin (Thermo Fisher Scientific) according to the manufacturer’s instruction. Briefly, 150 pmol of biotinylated proteins were incubated with 50 µL of the beads for 30 mins to 1 hour at room temperature with gentle rotation. The beads were then washed with TBST (20 mM Tris-HCl at pH 7.5, 100mM NaCl, and 0.025% Tween-20) five times using the DynaMag-2 magnetic holder (Thermo Fisher Scientific) and then resuspended to the original volume.

#### Antibody selection against HAs

Selection of antibodies against HA constructs were carried out in parallel. Briefly, the pre-selection library was mixed with 25 µL of beads coated with each HAs and incubated for 1 hour at room temperature with gentle rotation. After incubation, the beads were washed thrice with 400 µL TBST. The beads were then resuspended in water. These samples were referred as the “post-selection libraries”.

#### Antibody selection against HAs with CR9114 competition

The CR9114 competition screen was performed as describe above with the addition of a CR9114 blocking step prior to the selection. In brief, 25 µL beads coated with HAs were blocked with 2uM CR9114 IgG^10^, followed by the addition of 7.5uL of input library and incubation for 1 hour at room temperature with gentle rotation. After incubation, the beads were washed thrice with 400 µL TBST. The beads were then resuspended in water. These samples were referred as the “post-selection libraries”.

### Next-generation sequencing of the scFv library

The pre-selection libraries, post-selection libraries, and the small pool assemblies selected for quality control were amplified using PrimeSTAR Max DNA Polymerase (Takara Bio) per manufacturer’s instruction with the following primers (5’-GTA AAA CGA CGG CCA GTT TCA GGG GAC AAT TAC TAA AGG AGT ATC C-3’ and 5’-CAG GAA ACA GCT ATG ACC CAC TCG TCA TCC TTG TAA TCG GAT CCT CCG GA-3’. The PCR product was purified using a Monarch Gel Extraction Kit (New England Biolabs). A second round of PCR was carried out to add the adapter sequence and index to the amplicons (**Table S9**). The final PCR products were sequenced on one SMRTcell 8M on a PacBio Revio system using the CCS sequencing mode and a 30-hour movie time.

### Analysis of next-generation sequencing data

Circular consensus sequences (CCSs) were generated from the raw subreads using SMRTLink v13.0, setting the parameters to require 99.9% accuracy and a minimum of 3 passes. CSSs in FASTQ format were parsed using the SeqIO module in BioPython^59^ and filtered based on the base calling quality score, where any read with more than five nucleotides of phred quality score <40 were removed. The adapter sequences were then identified on each read and trimmed from the scFv sequences. Reads that did not have the complete adapter sequences were also removed. The filtered reads were then aligned to the reference scFv sequences and classified into three categories: 1) natively paired scFvs with no mutation, 2) natively paired scFvs with mutation, 3) others (non-natively paired scFvs and incomplete assemblies). Only the reads encoding natively paired scFvs with no mutation were used for downstream analysis. Frequency (*F*) of a scFv *i* a given replicate *k* of a given antigen *s* was computed for each replicate as follows:

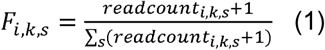

A pseudocount of 1 was added to each mutant to avoid division by zero in subsequent steps. We then calculated the enrichment (*E*) of a scFv *i* of a given replicate *k* of a given antigen *s* after the mRNA display selection as follows:

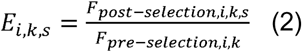

We calculated the mean enrichment of a scFv *i* of a given antigen *s* over two replicates, then inferred the binding score (*BS*) of a scFv *i* of a given antigen *s* using robust scaling:

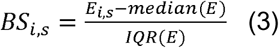

Where *median(E)* represents the median of a given dataset, *IQR(E)* representing the interquartile range of a given dataset. Custom cutoffs were set for each screen to determine hits (**Table S10**). For a given antigen *s*, the CR9114 competition index (*CI*) of a scFv *i* was computed as below:

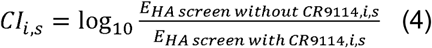

### Expression and purification of Fabs and IgGs

Heavy and light chains of the antibodies were cloned into a phCMV3 vector with a mouse immunoglobulin kappa signal peptide in human IgG1 Fc or Fab format. Plasmids encoding the heavy and light chains of antibodies were transfected into Expi293F cells using an ExpiFectamine 293 transfection kit (Gibco) in a 2:1 mass ratio for IgG or a 1:1 mass ratio for Fab following the manufacturer’s protocol. Supernatant was harvested at 6 days post-transfection and centrifuged at 4,000 xg for 30 mins at 4°C to remove cells and debris. The supernatant was subsequently clarified using a polyethersulfone membrane filter with a 0.22 mm pore size (Millipore). Antibodies were first purified by CaptureSelect CH1-XL beads (Thermo Fisher Scientific). Then, the antibodies were further purified by size exclusion chromatography using a HiLoad 16/100 Superdex 200 prep grade column (Cytiva) in 1x PBS. The A280 were measured using the Nanodrop One (Thermo Fisher Scientific) to calculate the sample concentration. Antibodies were stored at 4°C until used.

### Expression and purification of FLAG-tagged CR9114 scFv

CR9114 scFv nucleotide sequence with a pelB secretion peptide at the N-terminal and a FLAG-tag (DYKDDDK) followed by a stop codon at the C-terminal were synthesized (Integrated DNA Technologies) and ligated into a pET-28a plasmid vector backbone using NEBuilder HiFi DNA Assembly Master Mix (New England Biolabs). The ligated product was then transformed into DH5α competent cells (Thermo Fisher Scientific), and the plasmids were extracted using a QIAprep Spin Miniprep Kit (Qiagen). The plasmids were then transformed into BL21(DE3) competent cells (Thermo Fisher Scientific), followed by overnight growth at 37°C, 225 rpm shaking. 100 mL LB culture (Fisher Scientific) were inoculated at 1:500 ratio using the overnight culture and shaken at 37°C, 225 rpm. Once the OD_600_ reached approximately 0.6, IPTG (1 mM final concentration) was added to induce protein expression for 16 hours at 30°C, 200 rpm.

The overnight culture was centrifuged at 4,500 xg, 4°C for 1 hour to remove the supernatant. The pellet was resuspended using 2 mL of ice-cold 1x TES buffer (200 mM Tris-HCl at pH 8.0, 0.65 mM EDTA, and 0.5 M sucrose). The resuspended mixture was incubated at 4°C for 2 hours with gentle shaking. 5 mL of ice-cold 0.25x TES buffer was then added, followed by incubation at 4°C overnight. The mixture was centrifuged at 4,500 xg, 4°C for 1 hour to remove the pellet. The supernatant was subsequently clarified using a polyethersulfone membrane filter with a 0.22 mm pore size (Millipore) and purified using ANTI-FLAG M2 Affinity Gel (Millipore-Sigma) per manufacturer’s instructions. The A280 was measured using the Nanodrop One (Thermo Fisher) to calculate the sample concentration.

### Expression, purification, and quantitation of selected scFvs for validation

Nucleotide sequences of selected scFvs with a T7 promoter at the N-terminal and a FLAG-tag (DYKDDDK) followed by a stop codon at the C-terminal were synthesized (Integrated DNA Technologies) and amplified by PCR using Prime STAR Max DNA polymerase (Takara Bio). The PCR product was purified using a PureLink PCR purification kit (Thermo Fisher Scientific) and used as the template for *in vitro* translation using a PURExpress In Vitro Protein Synthesis Kit (New England Biolabs) with the addition of PURExpress Disulfide Bond Enhancer (New England Biolabs). The translated scFvs were then reverse purified using Pierce High-Capacity Ni-IMAC magnetic beads (Thermo Fisher Scientific) per manufacturer’s instructions to remove all His-tagged translation kit components.

To measure the concentration of the purified scFv, quantitation assays were performed by biolayer interferometry (BLI) using an Octet Red instrument (Sartorius). Briefly, rat anti-FLAG tag monoclonal antibody (L5) (Thermo Fisher Scientific) at 5 μg/mL in 1x kinetics buffer (1x PBS at pH 7.4, and 0.002% v/v Tween 20) were loaded onto ProG biosensors (Sartorius), then incubated with the 40x diluted scFv sample (5 μL of sample added to 195 μL of 1x kinetics buffer). The standard curve was generated via 2-fold serial dilutions using FLAG-tagged CR9114 scFv. The assay consisted of five steps: (1) baseline: 60 s with 1x kinetics buffer; (2) antibody capture: 180 s with rat anti-FLAG antibody; (3) baseline: 60 s with 1x kinetics buffer; (4) binding rate measurement: 120 s with standard and scFv samples; and (5) regeneration: 5s in regeneration buffer (0.1 M Glycine at pH 3.0) followed by 5 s in neutralization buffer (1 M Tris-HCl at pH 7.5), repeated 3 times. The data were analyzed using Octet analysis software 9.0, where the first 30 s of the binding rate measurement were used for final quantitation.

### Validation of oPool^+^ display via biolayer interferometry

The binding assay was performed by biolayer interferometry (BLI) using an Octet Red instrument (Sartorius). 1x kinetics buffer (1x PBS at pH 7.4, and 0.002% v/v Tween 20) were used for all experiments. Details of each experiment were described below:

#### Systematic binding validation

SA biosensors and Fab2G biosensors (Sartorius) were used for to validate binding of selected antibodies against the nine HA constructs. HA or Fab constructs at 20 μg/mL in 1x kinetics buffer were loaded onto the biosensors and incubated with 10 μg/mL Fabs/HAs. Measurements with NA and each Fab were also taken to serve as the baseline. The assay consisted of five steps: (1) baseline: 60 s with 1x kinetics buffer; (2) loading: 60 s with biotinylated HA or Fab; (3) baseline: 60 s with 1x kinetics buffer; (4) association: 60 s with Fab or HA samples; and (5) dissociation: 60 s with 1x kinetics buffer. The binding response during the association step were recorded.

#### Fab K_D_ measurement

Biotinylated H1 or H3 stem construct at 0.5 μM in 1x kinetics buffer was loaded onto SA biosensors (Sartorius) and incubated with 33 nM, 100 nM, and 300 nM of purified Fabs. The assay consisted of five steps: (1) baseline: 60 s with 1x kinetics buffer; (2) loading: 120 s with biotinylated HA stem domains; (3) baseline: 60 s with 1x kinetics buffer; (4) association: 120 s with Fab samples; and (5) dissociation: 120 s with 1x kinetics buffer. For estimating the exact K_D_, a 1:1 binding model was used.

#### scFv K_D_ measurement

Biotinylated H1 or H3 stem construct at 0.5 μM in 1x kinetics buffer was loaded onto SA biosensors (Sartorius) and incubated with 9x dilution (20 μL of sample added to 160uL of 1x kinetics buffer) and 18x dilution (10 μL of sample added to 170 μL of 1x kinetics buffer) of the purified scFv sample. The assay consisted of five steps: (1) baseline: 60 s with 1x kinetics buffer; (2) loading: 180 s with biotinylated HA stem domains; (3) baseline: 60 s with 1x kinetics buffer; (4) association: 120 s with Fab samples; and (5) dissociation: 120 s with 1x kinetics buffer. For estimating the exact K_D_, a 1:1 binding model was used.

#### CR9114 competition assay

SA biosensors and NTA biosensors (Sartorius) were used for to validate CR9114 competition of selected antibodies. HA constructs at 20 μg/mL in 1x kinetics buffer were first loaded onto the biosensors, then incubated with 20 μg/mL CR9114 Fab until saturation in binding, immediately followed by incubation with the selected Fabs. A no-CR9114 control experiment was performed concurrently for each Fab-antigen pair. The assay consisted of five steps: (1) baseline: 60 s with 1x kinetics buffer; (2) loading: 480 s with HAs; (3) baseline: 60 s with 1x kinetics buffer; (4) first association: 180 s with CR9114 Fab or 1x kinetics buffer; and (5) second association: 120 s with 10 μg/mL selected Fabs. Of note, competition assays of SI06 with 009-10-1G06, 045-09-1G05, 009-10-2F01, and 051-10-2B05 were done in the reverse order, where the first association consist of 900 s incubation with the selected Fabs, followed by 120 s second association with 10 μg/mL CR9114 Fab. The binding response of the first 60 s in step (5) were recorded, and the CR9114 competition % of a given antibody *i* against antigen *s* were calculated as below:

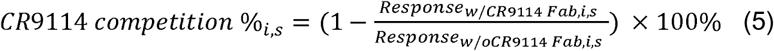

### Expression and purification of HA ectodomains

The HA ectodomains of H1N1 A/Puerto Rico/8/1934, H1N1 A/Beijing/262/1995, H1N1 A/Brisbane/02/2018 were cloned, expressed, and purified as mentioned above for the HA constructs used for screening. The HA ectodomains of H1N1 A/California/04/2009 (NR-15749), H5N1 A/bald eagle/Florida/W22-134-OP/2022 (NR-59476), H5N2 A/snow goose/Missouri/CC15-84A/2015 (NR-50651), and H5N8 A/northern pintail/WA/40964/2014 (NR-50174) were obtained from BEI Resources (https://www.beiresources.org/). The HA ectodomains of H1N1 A/USSR/90/1977 and H1N1 A/Taiwan/01/1986 were purchased from SinoBiological.

### Cryo-EM sample preparation, data collection, and data processing

#### AG11-2F01 Fab in complex with H1/SI06 HA

The AG11-2F01 Fab was incubated with H1/SI06 HA on ice overnight followed by size exclusion chromatography. The peak fraction of the Fab-HA complex was concentrated to around 1 mg/mL for cryo-EM sample preparation. Cryo-EM grids were prepared using a Vitrobot Mark IV (Thermo Fisher Scientific). 3.5 μL of the sample was applied to a 300-mesh Quantifoil R1.2/1.3 Cu grid pretreated with glow-discharge. Excess liquid was blotted away using filter paper with blotting force -5 and blotting time 3 s. The grid was then flash frozen in liquid ethane. Movies were then collected on a Titan Krios microscope equipped with Gatan BioQuantum K3 imaging filter and camera (Thermo Fisher Scientific). Images were recorded at 130,000x magnification, corresponding to a pixel size of 0.33 Å/pixel at super-resolution mode of the camera. A defocus range of -0.8 µm to -1.5 µm was set. A total dose of 50 e^-^/Å^2^ of each exposure was fractionated into 50 frames. Both untilted and 30-degree-tilted data were collected and combined to alleviate the preferred orientation problem of the sample.

CryoSPARC^60^ was used to process the cryo-EM data. For model building, ABodyBuilder^61^ was used to generate an initial model for AG11-2F01 Fab. This model, together with the model of H1/SI06 HA (PDB 6FYT)^62^, was fitted into the cryo-EM density map using UCSF Chimera^63^. The model was manually adjusted in Coot^64^ and refined with Phenix real-space refinement program^65^. This process was iterated for several cycles until no significant improvement of the model was observed.

#### 16.ND.92 Fab in complex with H1/SI06 HA

The 16.ND.92 Fab was incubated with H1/SI06 HA and FISW84 Fab, a known HA anchor antibody^66^, on ice overnight followed by size exclusion chromatography. The peak fraction of the Fab-HA complex was concentrated to around 3 mg/mL for cryo-EM sample preparation. 0.1% (w/v) of n-octyl-ß-D-glucoside was added to reduce orientation bias. Cryo-EM grids were prepared using a Vitrobot Mark IV (Thermo Fisher Scientific). 3 μL of the sample was applied to a 400-mesh Quantifoil R1.2/1.3 Cu grid pretreated with glow-discharge. Excess liquid was blotted away using filter paper with blotting force 0 and blotting time 3 s. The grid was then flash frozen in liquid ethane. Movies were then collected on a Titan Krios microscope equipped with Gatan BioQuantum K3 imaging filter and camera (Thermo Fisher Scientific). Images were recorded at 81,000x magnification, corresponding to a pixel size of 0.53 Å/pixel at super-resolution mode of the camera. A defocus range of -0.5 µm to -5 µm was set. A total dose of 57.35 e^-^/Å^2^ of each exposure was fractionated into 40 frames.

CryoSPARC^60^ was used to process the cryo-EM data. DeepEMhancer^67^ was used to generate the sharpened density map for downstream model building. For model building, IgFold^68^ was used to generate an initial model for 16.ND.92 Fab. This model, together with the model of H1/SI06 HA (PDB 6FYT)^62^, was fitted into the cryo-EM density map using Phenix DockinMap module^69^. The models were manually adjusted in Coot^64^ and refined with Phenix real-space refinement program^65^. This process was iterated for several cycles until no significant improvement of the model was observed.

### Structural analysis of HA-antibody complexes

Buried surface areas upon binding and paratope residues of AG11-2F01, 16.ND.92, MEDI8852 (PDB 5JW4), 56.a.09 (PDB 5K9J), 54-1G05 (PDB 6WIZ), PN-SIA28 (PDB 8GV5), 39.29 (PDB 4KVN), and 429 B01 (PDB 6NZ7)^7,9,31–34^ were analyzed using PDBePISA^70^. The CDRH3 region and IGHD 3-3 usage of each antibody was annotated using IgBLAST^71^. The molecular interactions of AG11-2F01 and 16.ND.92 in complex with H1/SI06 HA were analyzed and visualized using PyMOL (Schrödinger).

### Enzyme-linked immunosorbent assay (ELISA)

Nunc MaxiSorp ELISA plates (Thermo Fisher Scientific) were coated overnight at 4°C with 100 μL of recombinant proteins at 1 μg/mL in 1x PBS. The next day, plates were washed thrice with 1x PBS containing 0.05% Tween 20 and blocked with 200 μl of 5% non-fat milk in 1x PBS for 2 hours at room temperature. For systematic validation of oPool^+^ display, 10 μg/mL of monoclonal antibodies were added to the plates, and incubated for 2 hours at 37°C; for functional characterization of AG11-2F01 and 16.ND.92, monoclonal antibodies were serially diluted 10-fold starting from 100 μg/mL, added to the plates, and incubated for 2 hours at 37°C. Plates were then washed thrice and incubated with horseradish peroxidase (HRP)-conjugated goat anti-human IgG antibody (Thermo Fisher Scientific) at 1:5,000 dilution for 1 hour at 37°C. After six washes with 1x PBS containing 0.05% Tween 20, 100 μL of 1-Step TMB ELISA Substrate Solution (Thermo Fisher Scientific) was added to each well. After incubation for 10 mins, the reaction was stopped with 50 μL of 2 M H_2_SO_4_ solution, and absorbance values were measured at 450 nm (OD_450_) using a BioTek Synergy HTX Multimode Reader (Agilent).

### Recombinant virus construction and purification

Recombinant viruses with HA and NA segments from the indicated H1N1 strains and six internal segments from H1N1 A/Puerto Rico/8/1934 (PR8) were obtained from BEI Resources (https://www.beiresources.org/). Recombinant viruses were rescued using the eight-plasmid reverse genetics system^72^. Briefly, plasmids encoding the HA segments from H1N1 A/California/07/2009 and H1N1 A/Michigan/45/2015 along with seven plasmids encoding the other seven segments from PR8 were transfected into a co-culture of HEK293T (human embryonic kidney) cells and MDCK-SIAT1 (Madin-Darby Canine Kidney) cells at a 6:1 ratio. Supernatants were injected into 8-10 days old embryonated chicken eggs and incubated at 37°C for 48 hours. Viruses in the allantoic fluid were plaque-purified on MDCK-SIAT1 cells grown in Dulbecco’s Modified Eagles Medium (Gibco) containing 10% fetal bovine serum (Gibco) and a penicillin-streptomycin mix (100 U/mL penicillin and 100 μg/mL streptomycin, Gibco). The HA sequence of each virus was confirmed by Sanger sequencing.

### Microneutralization assay

For the microneutralization assay, MDCK-SIAT1 cells were seeded in 96-well plates. After reaching 100% confluency, MDCK-SIAT1 cells were washed once with 1x PBS. Minimal essential media (Gibco) containing 25 mM HEPES (Gibco) was then added to the cells. Monoclonal antibodies were serially diluted 10-fold starting from 100 μg/ml and mixed with 100 TCID_50_ (median tissue culture infectious dose) of viruses at equal volume and incubated at 37°C for 1 hour. Subsequently, the mixture was inoculated into cells and incubated at 37°C for another hour. Cell supernatants were discarded and replaced with minimal essential media containing 25 mM HEPES, and 1 μg/mL TPCK-trypsin (Sigma). Plates were incubated at 37°C for 72 hours, and virus presence was detected by hemagglutination assay to determine the MN_50_ titers.

### Mice

The animal experiments were performed in accordance with protocols approved by UIUC Institutional Animal Care and Use Committee (IACUC). Six-week-old female BALB/c mice (Jackson Laboratory) were used for all animal experiments.

### Prophylactic and therapeutic protection experiments

Female BALB/c mice at 6 weeks old (n = 5 per group) were anesthetized with isoflurane and intranasally infected with 5x lethal dose (LD_50_) of recombinant PR8 virus. Mice were given the indicated antibody at a dose of 5 mg/kg intraperitoneally at 4 hours before infection (prophylaxis) or 4 hours after infection (therapeutics). Weight loss was monitored daily for 14 days. The humane endpoint was defined as a weight loss of 25% from initial weight at day 0. Of note, while our BALB/c mice were not modified to facilitate interaction with human IgG1, human IgG1 could interact with mouse Fc gamma receptors^73–75^. To determine the lung viral titers at day 3 post-infection, lungs of infected mice were harvested and homogenized in 1 mL of minimal essential media with 10% bovine serum albumin using a gentleMACS C Tube (Miltenyi Biotec). Subsequently, virus titers were measured by TCID_50_ assay.

## Supporting information

Table S1: Selected HA antibodies.

Table S2: Oligo pool sequences.

Table S3: Enrichment results of oPool+ display.

Table S4: Final screening results

## ACKNOWLEDGEMENTS

We thank Anders Olson, Chris Brooke, David Kranz, and Steve Sligar for helpful discussion and the Roy J. Carver Biotechnology Center at the University of Illinois at Urbana-Champaign for assistance with PacBio sequencing. We thank Kristen Flatt and the Materials Research Laboratory Central Research Facilities at the University of Illinois at Urbana-Champaign for access to cryo-EM instrumentation during the screening of the Fab-HA complexes. We are grateful to the Cryo-EM Core and the Core Facility for Advanced Research Computing at Case Western Reserve University for the access of cryo-EM instrumentation and data processing. We also thank Frank Vago and the Cryo-EM Facility at Purdue University for the access of cryo-EM instrumentation and data collection. This work is supported by National Institutes of Health R01 AI167910 (N.C.W.), DP2 AT011966 (N.C.W.), the Searle Scholars Program (N.C.W.), Howard Hughes Medical Institute Emerging Pathogens Initiative (N.C.W.), UIUC William T. and Lynn Jackson Graduate Student Fellowship (W.O.O.), UIUC Gregorio Weber Graduate Student Fellowship (W.O.O.), and Carl R. Woese Institute for Genomic Biology Postdoctoral Fellowship (H.L.).

## AUTHOR CONTRIBUTIONS

W.O.O. and N.C.W. conceived and designed the study. W.O.O., H.L., W.L., Y.W., K.E.D. and N.C.W. developed the methodology. W.O.O. and L.T. assembled the library and performed mRNA display experiment. W.O.O., W.L. and N.C.W. analyzed the PacBio sequencing data. W.O.O., H.L., R.L., M.T., A.B.G. and L.A.R. expressed and purified recombinant proteins. W.O.O., Z.M., R.L., T.P., X.D., and N.C.W. performed structural analysis of the antibodies. W.O.O., H.L., D.C., and M.R.A performed functional characterization of the antibodies. X.D. and N.C.W. provided resources and support. W.O.O. H.L., W.L. and N.C.W. wrote the paper, and all authors reviewed and edited the paper.

## DECLARATION OF INTERESTS

N.C.W. consults for HeliXon. The authors declare no other competing interests.

## DATA & CODE AVAILABILITY

Raw sequencing data have been submitted to the NIH Short Read Archive under accession number: BioProject PRJNA1150188. Cryo-EM density maps and coordinates have been deposited to EMDB and PDB with accession numbers: EMD-46727 and EMD-45930; PDB 9DBX and 9CU7. Custom scripts as well as raw data for experimental validations have been deposited to: https://github.com/nicwulab/oPool_display.

## SUPPLEMENTAL FIGURES

Figure S1. Curation and design of the natively paired HA antibody library.

Figure S2. One-pot PCR assembly from 25 scFvs to 200 scFvs per tube.

Figure S3. Final library assembly and quality assessment.

Figure S4. HA stem domain and ectodomain HA screens.

Figure S5. Quality assessment of the HA stem and ectodomain HA screens. Figure S6. CR9114 competition screens.

Figure S7. Quality assessment of CR9114 competition screens.

Figure S8. BLI sensorgrams for systematic validation of oPool+ display.

Figure S9. BLI sensorgrams for stem antibody binding affinity measurements in Fab format.

Figure S10. BLI sensorgrams for stem antibody binding affinity measurements in scFv format.

Figure S11. Competition indexes of all antibody hits.

Figure S12. BLI sensorgrams for CR9114 competition assays.

Figure S13. CDR H3 sequence analysis of AG11-2F01 and 16.ND.92. Figure S14. ELISA titration curves.

## SUPPLEMENTAL TABLES

Table S1. Selected HA antibodies.

Table S2. Oligo pool sequences.

Table S3. Enrichment results of oPool+ display.

Table S4. List of antibody hits and their binding profiles.

Table S5. CR9114 competition data of validated antibodies

Table S6. Cryo-EM data collection, refinement and validation statistics.

Table S7. Buried surface areas upon binding of IGHD3-3 antibodies.

Table S8. Cost and time comparison between traditional approaches for antibody specificity characterization and oPool+ display.

Table S9. Sequences of primers used in this study.

Table S10. Custom cutoff for each screen.

## Notes

https://github.com/nicwulab/oPool_display

### Summary of Updates

In this lastest update, we have substantially expanded our screening effort with >5,000 binding tests performed. A more systematic and comprehensive binding validation was also adopted.

