## Supplementary material for "High-throughput synthesis and specificity characterization of natively paired antibodies using oPool^+^ display": Figure S1-S14 and Table S5-S10

**Table of Contents**

**SUPPLEMENTAL FIGURES**

Figure S1. Curation and design of the natively paired HA antibody library.

Figure S2. One-pot PCR assembly from 25 scFvs to 200 scFvs per tube.

Figure S3. Final library assembly and quality assessment.

Figure S4. HA stem domain and HA ectodomain screens.

Figure S5. Quality assessment of the HA stem and HA ectodomain screens.

Figure S6. CR9114 competition screens.

Figure S7. Quality assessment of CR9114 competition screens.

Figure S8. BLI sensorgrams for systematic validation of oPool^+^ display.

Figure S9. BLI sensorgrams for stem antibody binding affinity measurements in Fab format.

Figure S10. BLI sensorgrams for stem antibody binding affinity measurements in scFv format.

Figure S11. Competition indexes of all antibody hits.

Figure S12. BLI sensorgrams for CR9114 competition assays.

Figure S13. CDR H3 sequence analysis of AG11-2F01 and 16.ND.92.

Figure S14. ELISA titration curves.

**SUPPLEMENTAL TABLES**

Table S1. Selected HA antibodies.

Table S2. Oligo pool sequences.

Table S3. Enrichment results of oPool^+^ display.

Table S4. List of antibody hits and their binding profiles.

Table S5. CR9114 competition data of validated antibodies

Table S6. Cryo-EM data collection, refinement and validation statistics.

Table S7. Buried surface areas upon binding of IGHD3-3 antibodies.

Table S8. Cost and time comparison between traditional approaches for antibody specificity characterization and oPool^+^ display.

Table S9. Sequences of primers used in this study.

Table S10. Custom cutoff for each screen.

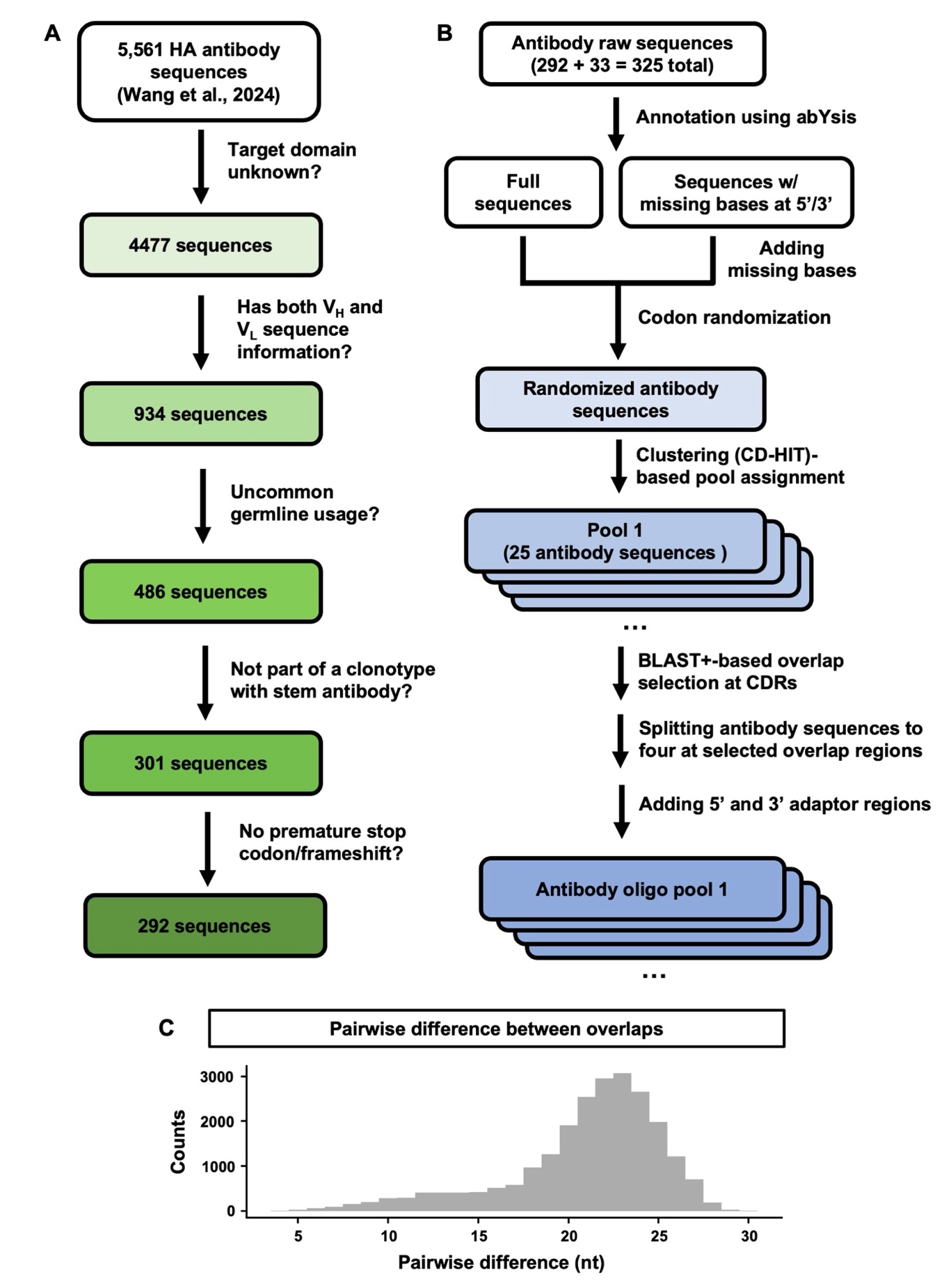

**Figure S1. Curation and design of the natively paired HA antibody library. (A)** Schematics of antibody selection for antibody library synthesis. The indicated criteria were applied to select antibodies from a previously curated influenza HA antibody database. **(B)** Schematics of our strategy for oligo pool design. **(C)** Distribution of the pairwise nucleotide differences across all overlaps.

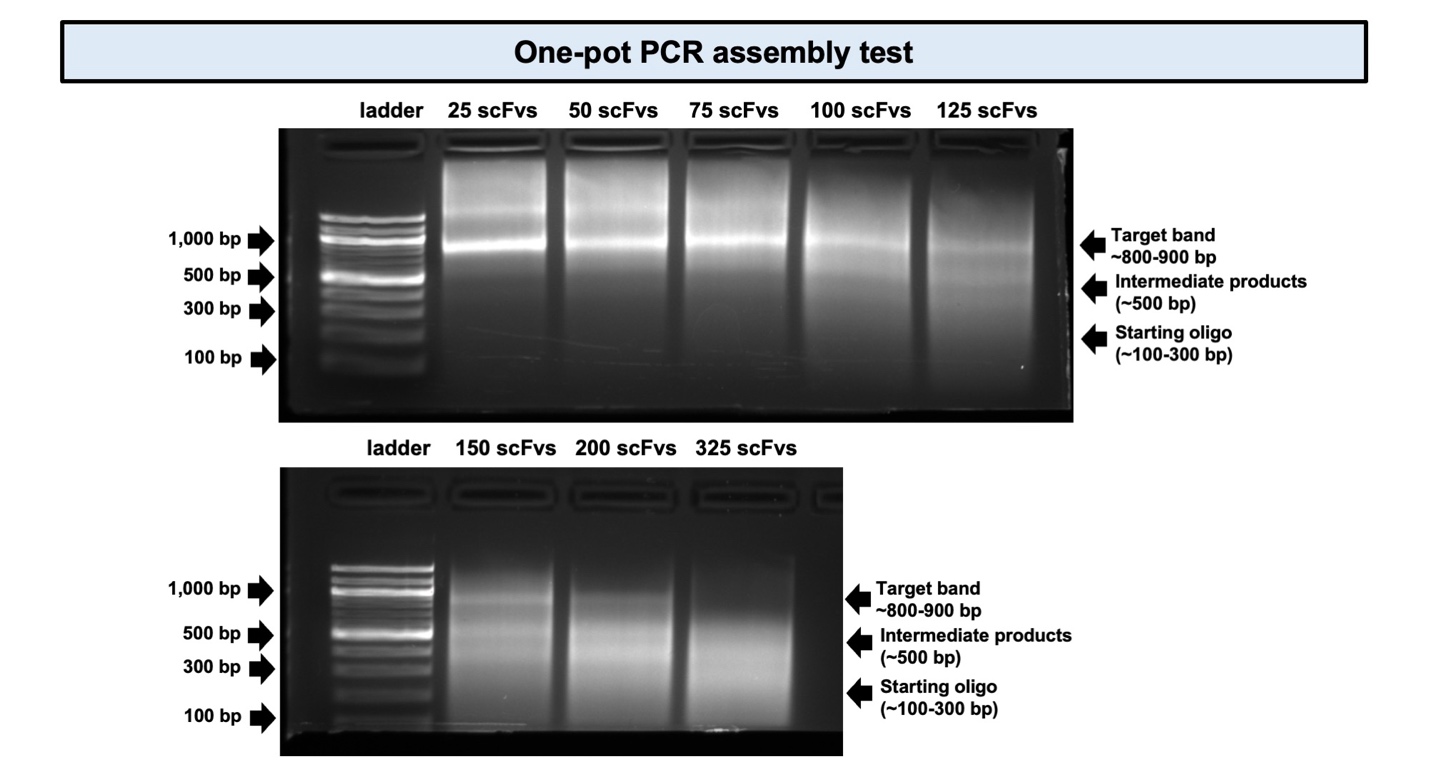

**Figure S2. One-pot PCR assembly from 25 scFvs to 325 scFvs per tube.** Agarose gel electrophoresis of antibody library assemblies from 25 scFvs per tube to 325 scFvs per tube. One of the two technical replicates are shown. Target bands (~800-900 bp), intermediate products (~500 bp), starting oligos (~100-300 bp), as well as the 1000 bp, 500 bp, 300bp, and 100 bp markers are indicated by arrows.

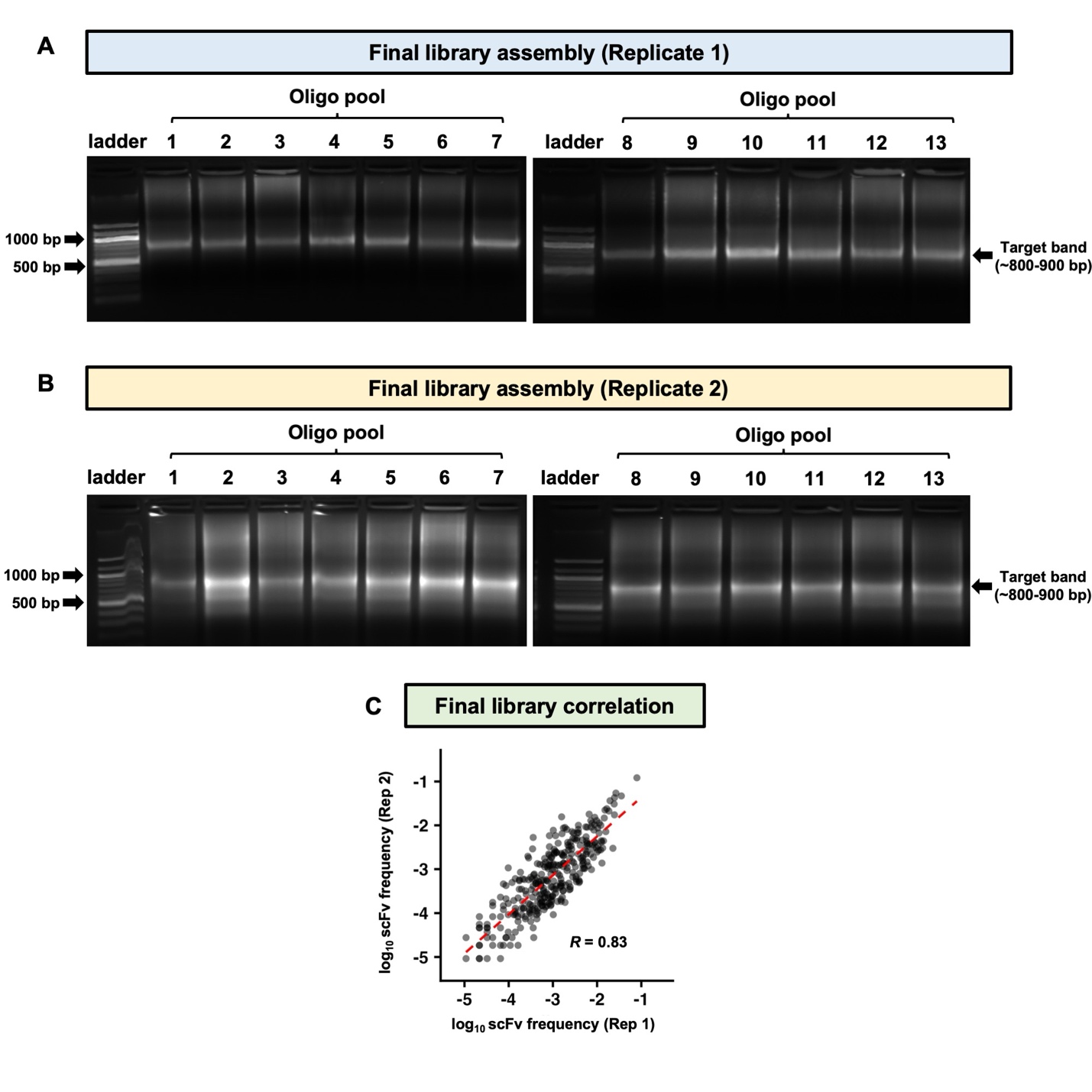

**Figure S3. Final library assembly and quality assessment. (A-B)** Agarose gel electrophoresis of antibody library assemblies from individual oligo pools with 25 scFvs each. Two technical replicates were performed. Target bands as well as the 1000 bp and 500 bp markers are indicated by arrows. **(C)** Correlations between replicates of final antibody library assemblies after equal molar pooling. Pearson correlation coefficient (*R*) between technical replicates is indicated.

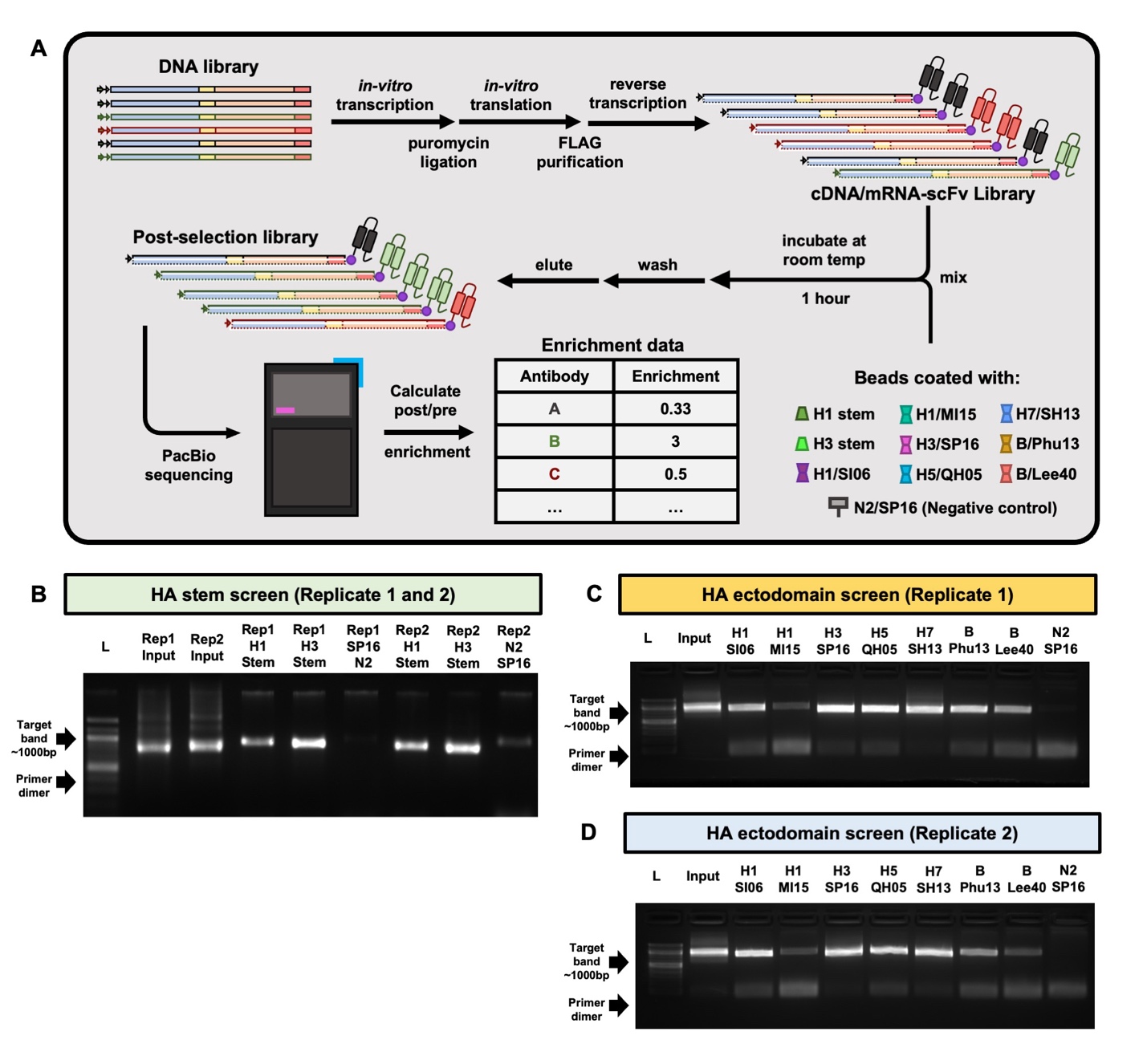

**Figure S4. HA stem domain and HA ectodomain screens. (A)** Schematic overview of the mRNA display selection. Different colors represent different scFvs. The purple dot represents the puromycin. H3N2 A/Singapore/INFIMH-16-0019/2016 NA was included as a negative control. **(B)** Agarose gel electrophoresis of pre and post selection library against H1 stem and H3 stem. **(C-D)** Agarose gel electrophoresis of pre and post selection library against the seven HA ectodomain constructs. Target bands and primer dimers are indicated by arrows.

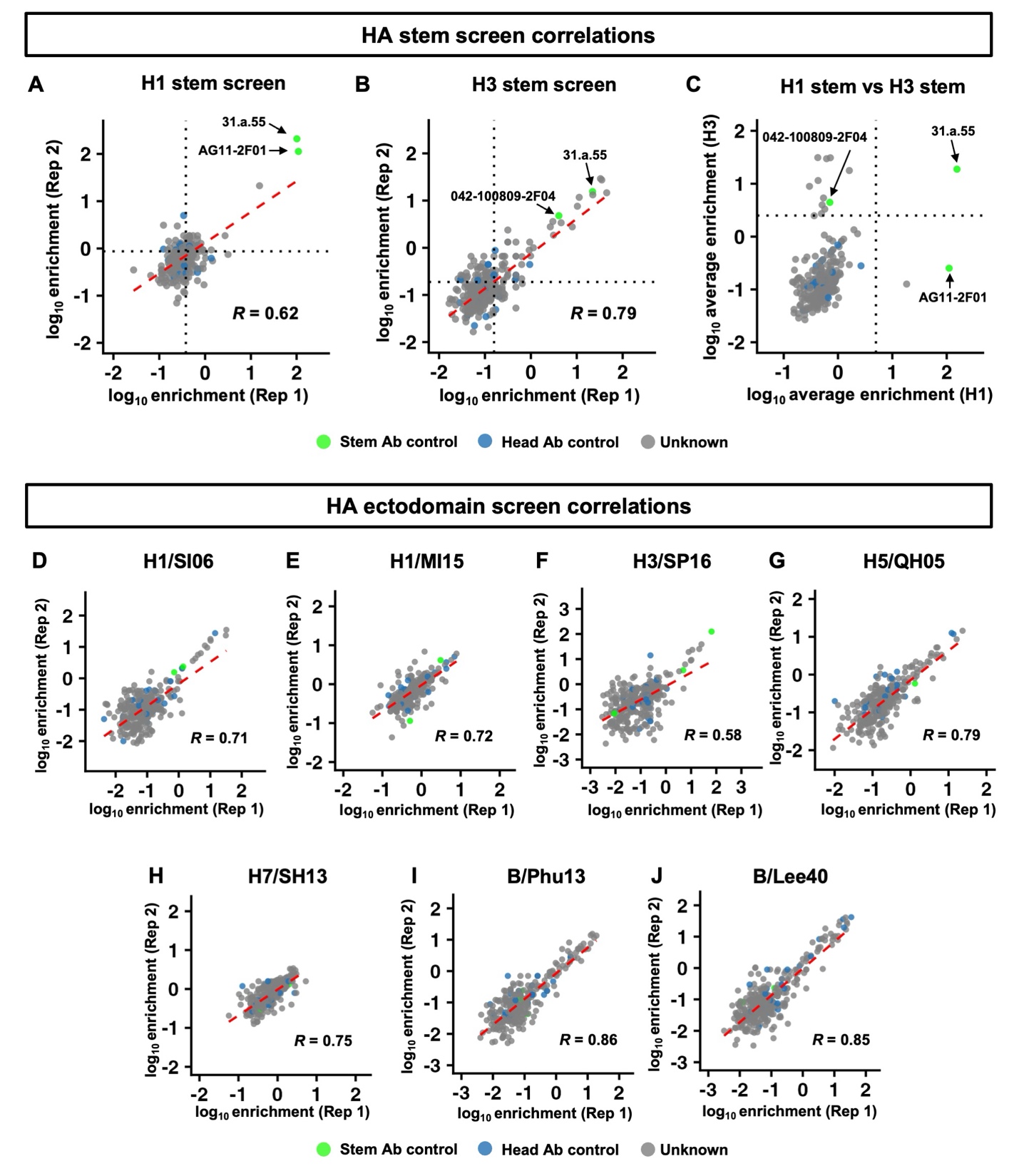

**Figure S5. Quality assessment of the HA stem and HA ectodomain screens.** **(A-B)** Correlation between replicates of mRNA display selection against H1 stem and H3 stem. For each replicate, the average enrichment of head antibody controls is shown as grey dotted lines. **(C)** The average enrichments against H1 stem and H3 stem are compared. The grey dotted lines represent the cutoff for the identification of HA stem antibody candidates. H1 stem antibody candidates are in lower right quadrant, whereas H3 stem antibody candidates are in the upper left quadrant. **(D-J)** Correlation between replicates of mRNA display selection against the seven HA ectodomain constructs. The green dots represent the known HA stem antibodies, whereas the blue dots represent known HA head antibodies. The three stem antibody controls in this study are 31.a.55 (both H1 and H3 stem-binding), AG11-2F01 (H1 stem-binding), and 042-100809-2F04 (H3 stem-binding) (*7*, *16*). Pearson correlation coefficient (*R*) between replicates is indicated. The linear fit lines are shown as red dash lines.

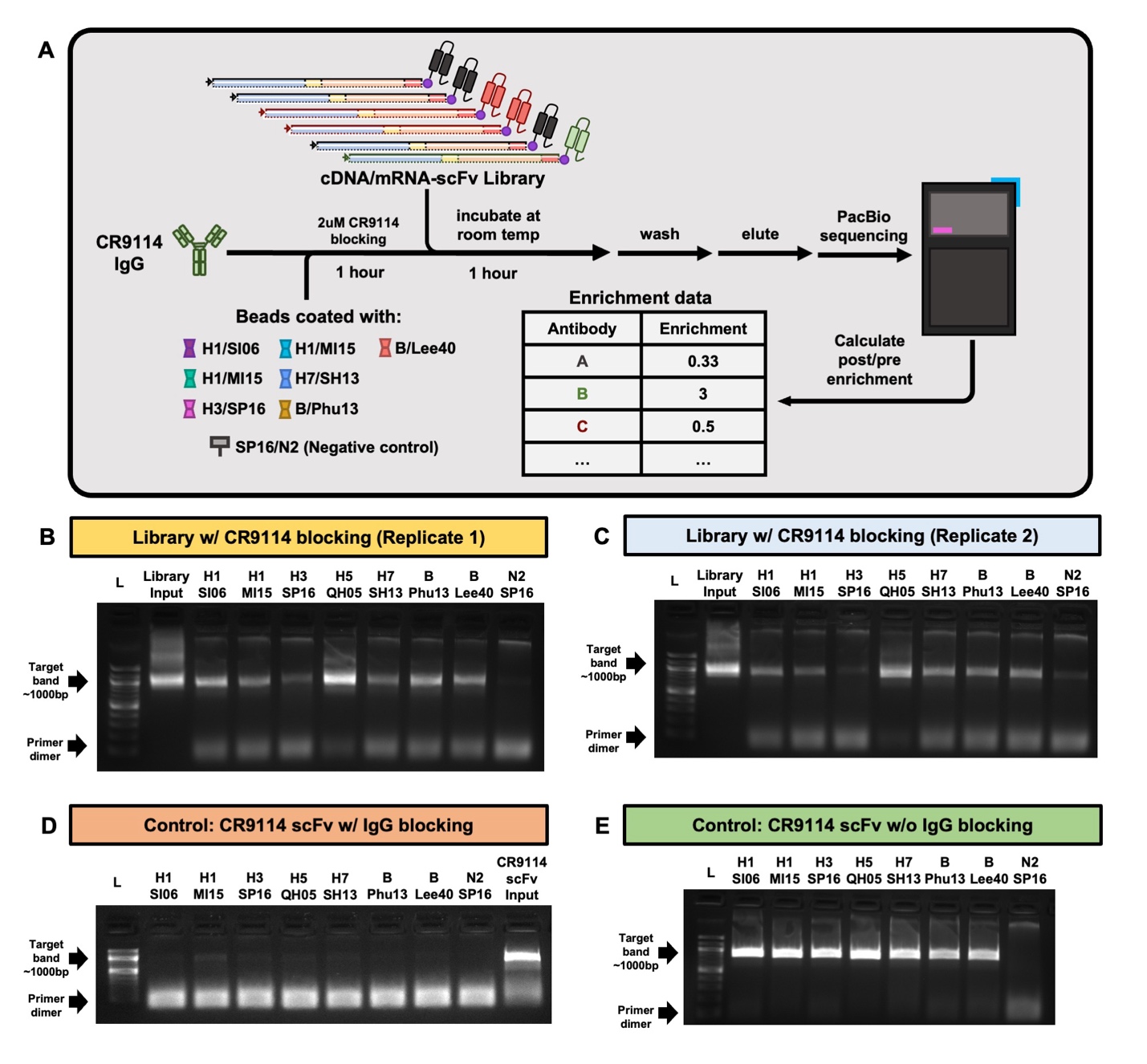

**Figure S6. CR9114 competition screens. (A)** Schematic overview of the mRNA display selection with CR9114 competition. Different colors represent different scFvs. The purple dot represents the puromycin. H3N2 A/Singapore/INFIMH-16-0019/2016 NA was included as a negative control. **(B-C)** Agarose gel electrophoresis of pre and post selection library against the seven HA ectodomain constructs with CR9114 competition. While H3/SP16 screen produced distinct band in replicate 1, the band intensity for H3/SP16 selection was similar to N2/SP16 in replicate 2. Therefore, the samples from the CR9114 competition screen for H3/SP16 were not sequenced. **(D-E)** Agarose gel electrophoresis of positive and negative controls pre- and post-selection with CR9114 scFv as input. Target bands and primer dimers are indicated by arrows.

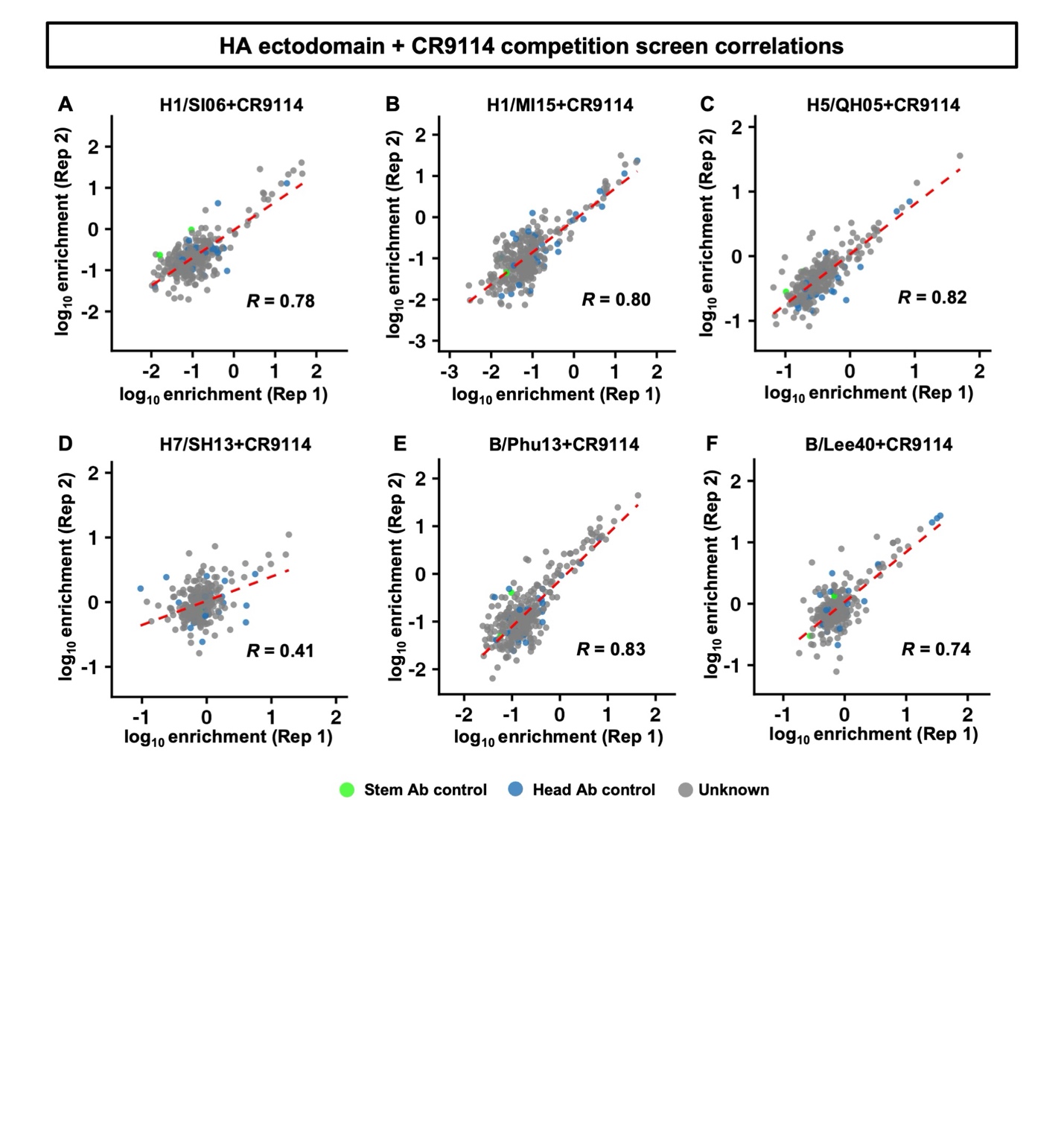

**Figure S7. Quality assessment of CR9114 competition screens.** Correlation between replicates of mRNA display selection against the seven HAs with CR9114 competition. Pearson correlation coefficient (*R*) between replicates is indicated. The linear fit line is shown as dotted red lines.

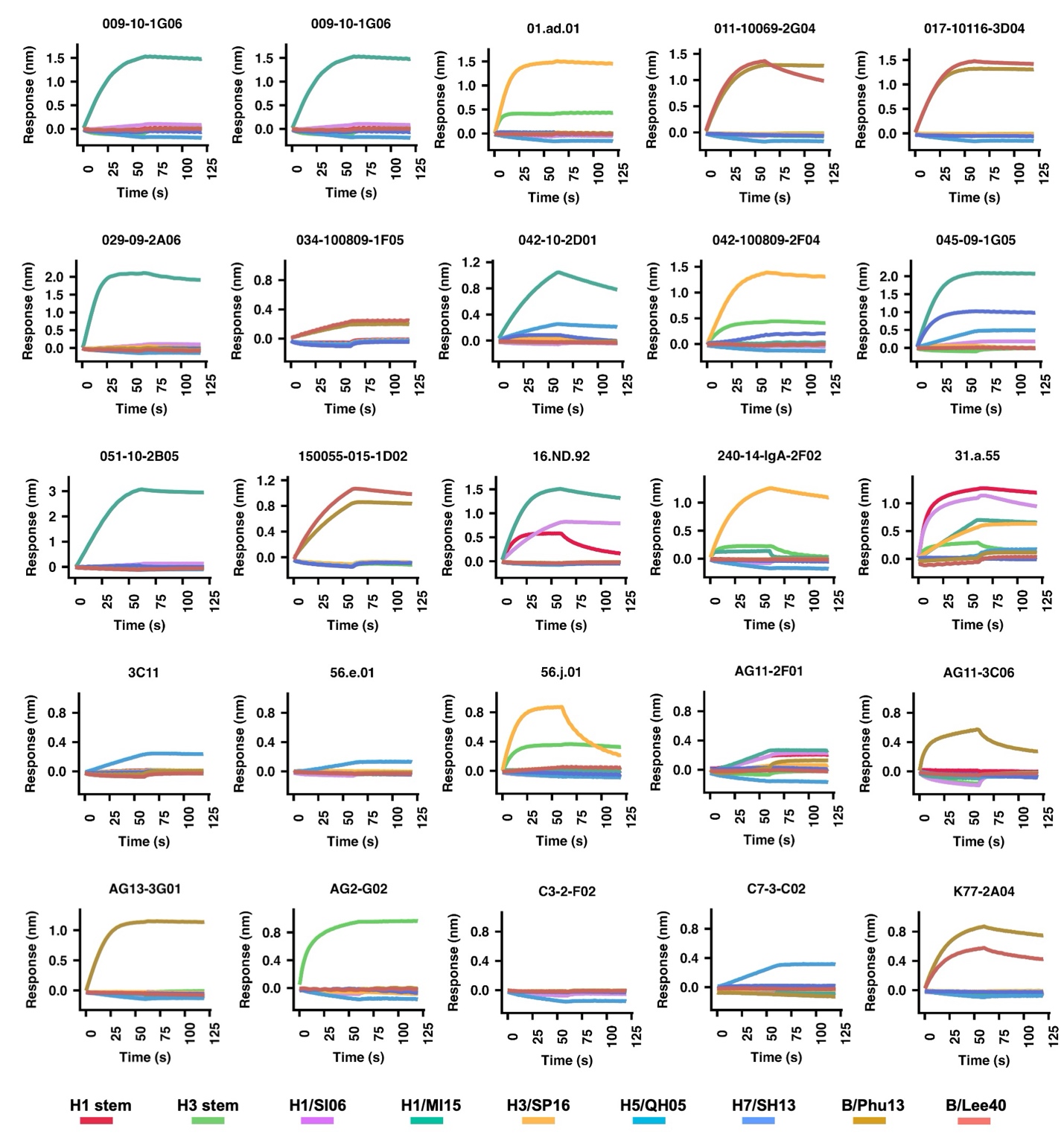

**Figure S8. BLI sensorgrams for systematic validation of oPool^+^ display.** The binding responses of the 25 selected antibody Fabs against all nine HA constructs used in oPool+ display. The binding profile of each antibody is shown in one sensorgram, with the antigens color coded.

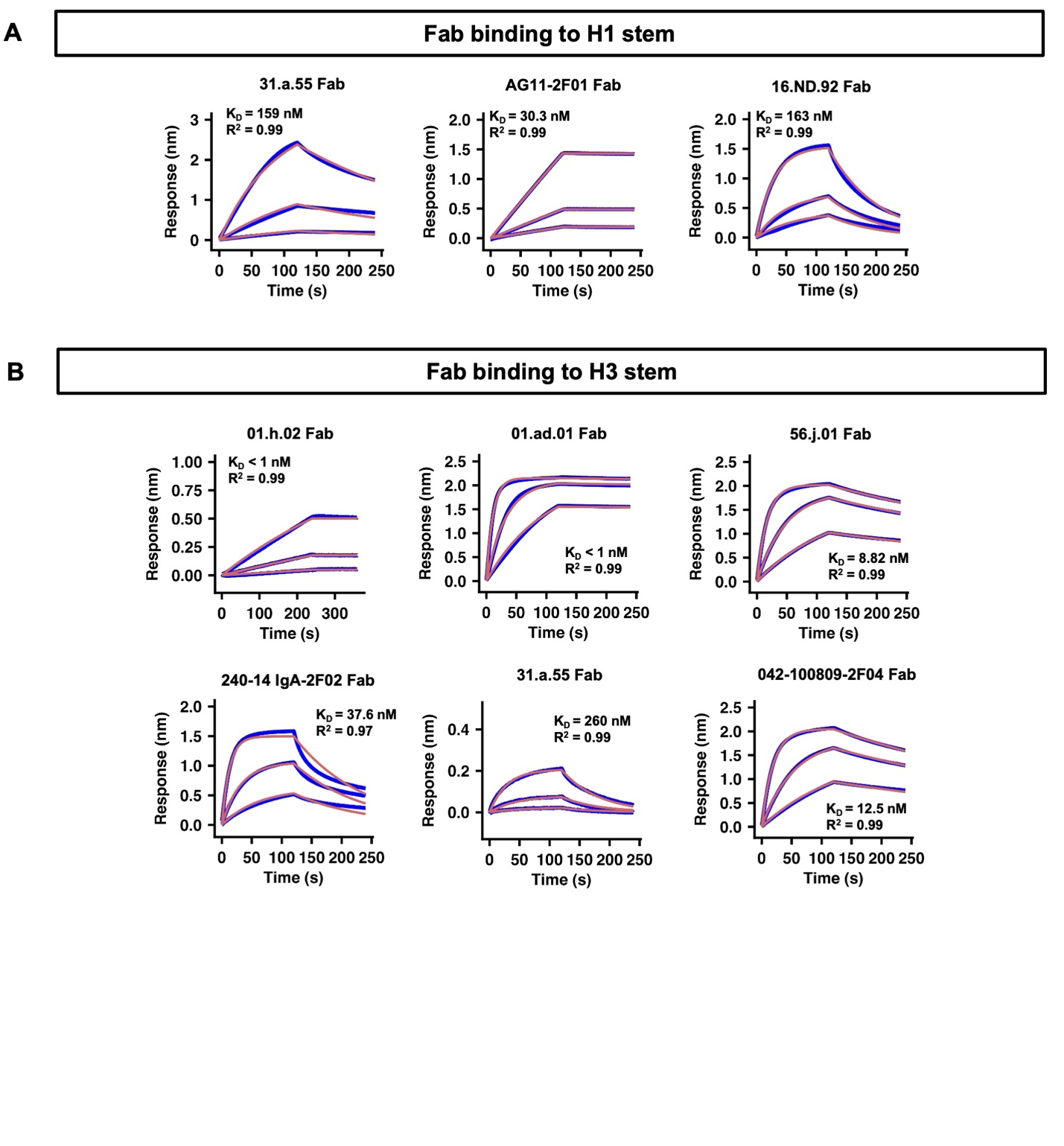

**Figure S9. BLI sensorgrams for antibody validation in Fab format.** Binding kinetics of selected antibodies as Fabs against **(A)** H1 stem and **(B)** H3 stem were measured by biolayer interferometry (BLI). Y-axis represents the response. Blue lines represent the response curves and red lines represent the best fit model (1:1 binding model). Binding kinetics were measured for three concentrations of Fab at 3-fold dilution ranging from 300 nM to 33 nM. Dissociation constants (K_D_) and goodness of fit values (R^2^) are indicated.

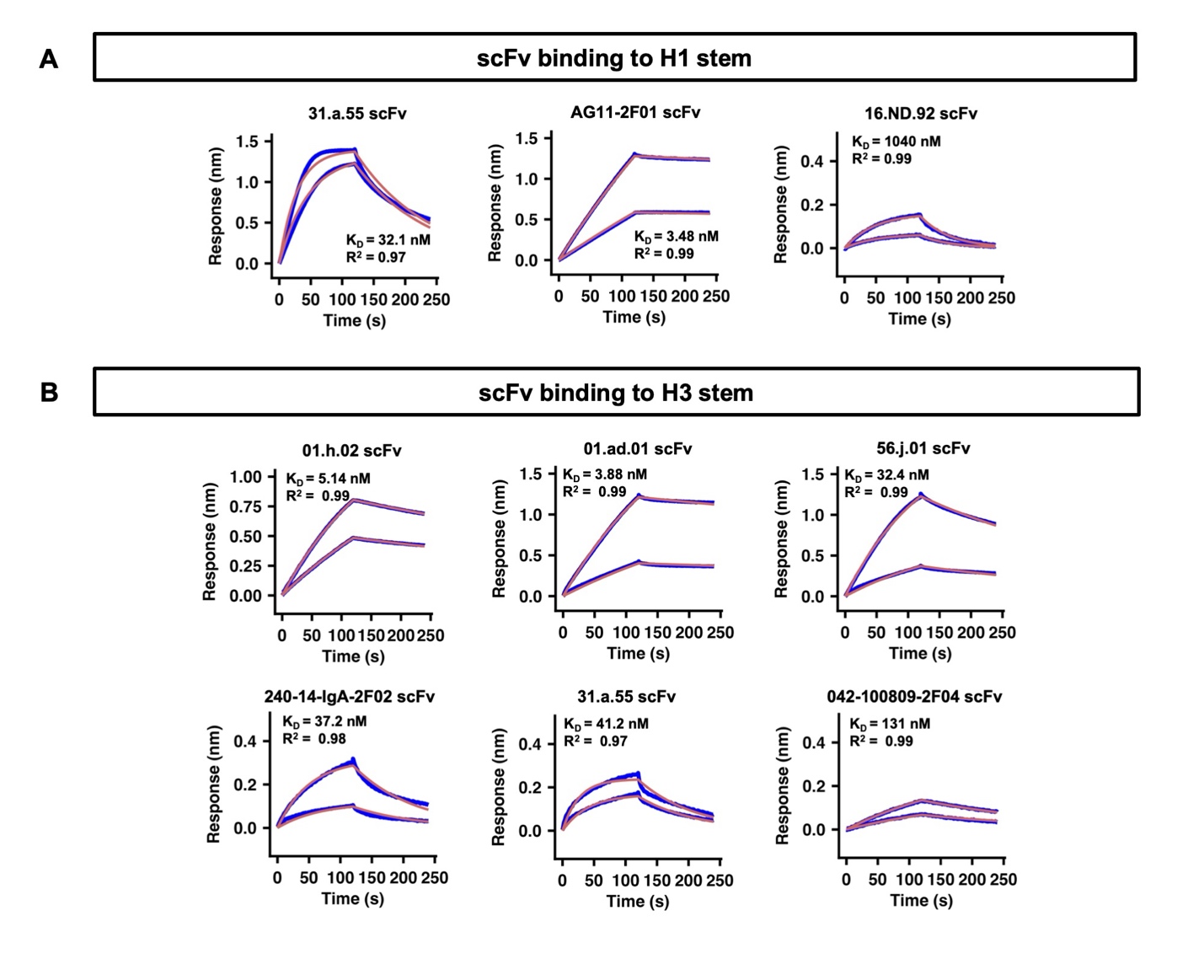

**Figure S10. BLI sensorgrams for antibody validation in scFv format. (A-B)** Binding kinetics of selected antibodies as scFvs against **(A)** H1 stem and **(B)** H3 stem were measured by biolayer interferometry (BLI). Y-axis represents the response. Blue lines represent the response curves and red lines represent the best fit model (1:1 binding model). Binding kinetics were measured for two concentrations of scFvs (9-fold and 18-fold dilution of the actual sample concentration). Dissociation constants (K_D_) and R^2^ are indicated.

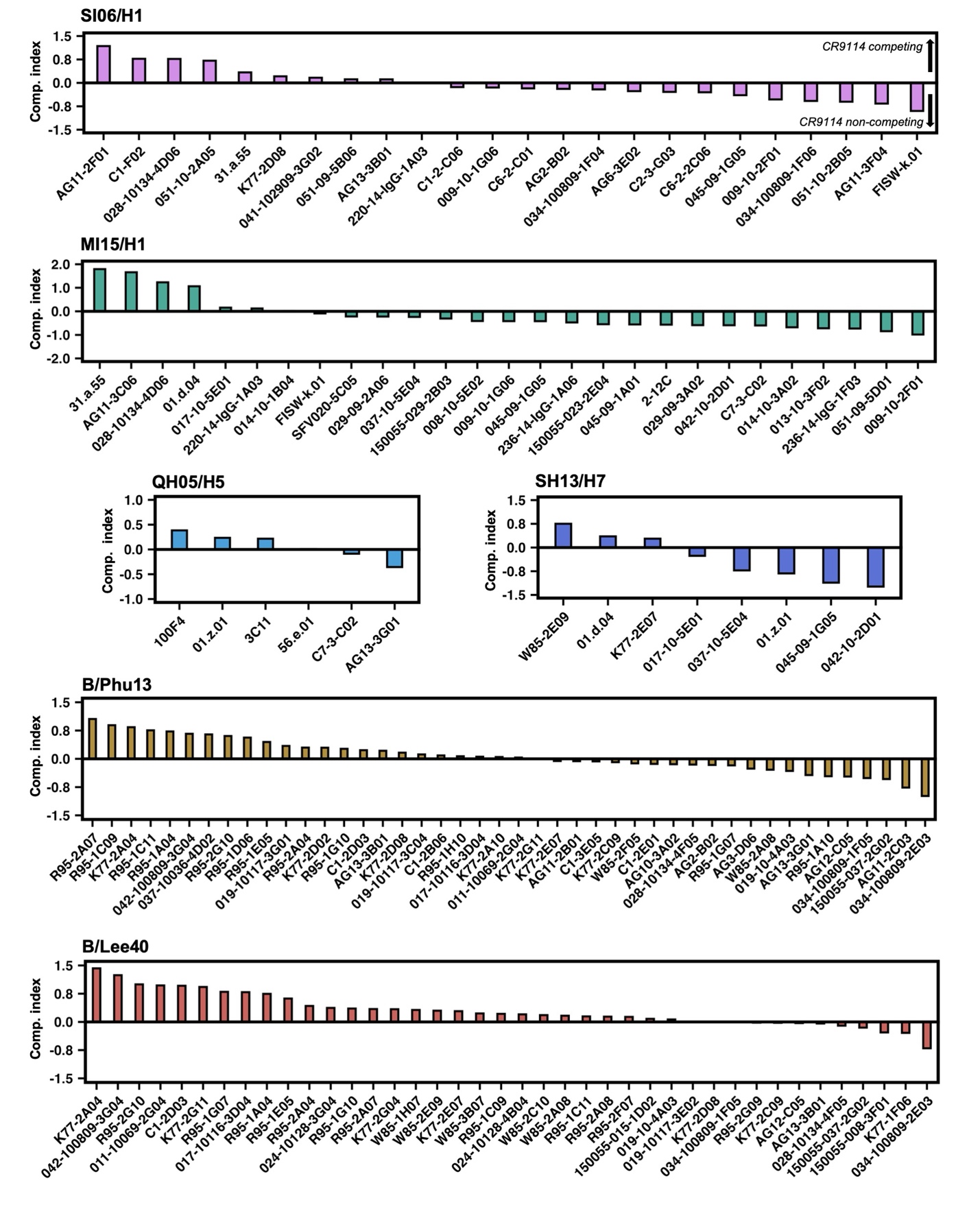

**Figure S11. Competition indexes of all antibody hits.** The CR9114 competition indices of all antibody hits. The competition indices calculated from oPool^+^ display were shown. High positive values indicate CR9114 competition, low positive and high negative values indicate no CR9114 competition.

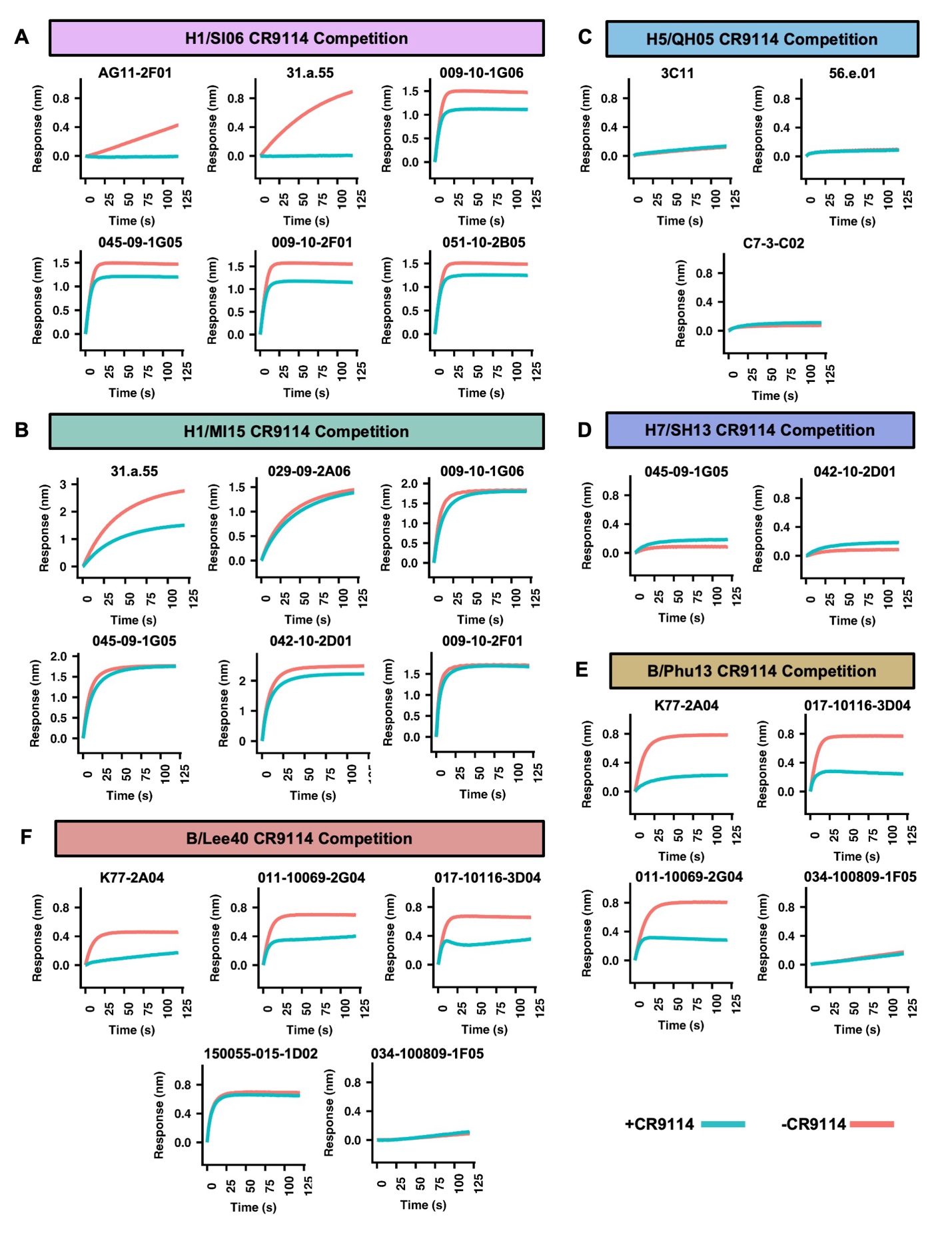

**Figure S12. BLI sensorgrams for CR9114 competition assays.** The binding responses of selected antigen-antibody pairs with (blue) or without (orange) prior association with CR9114 Fab are shown.

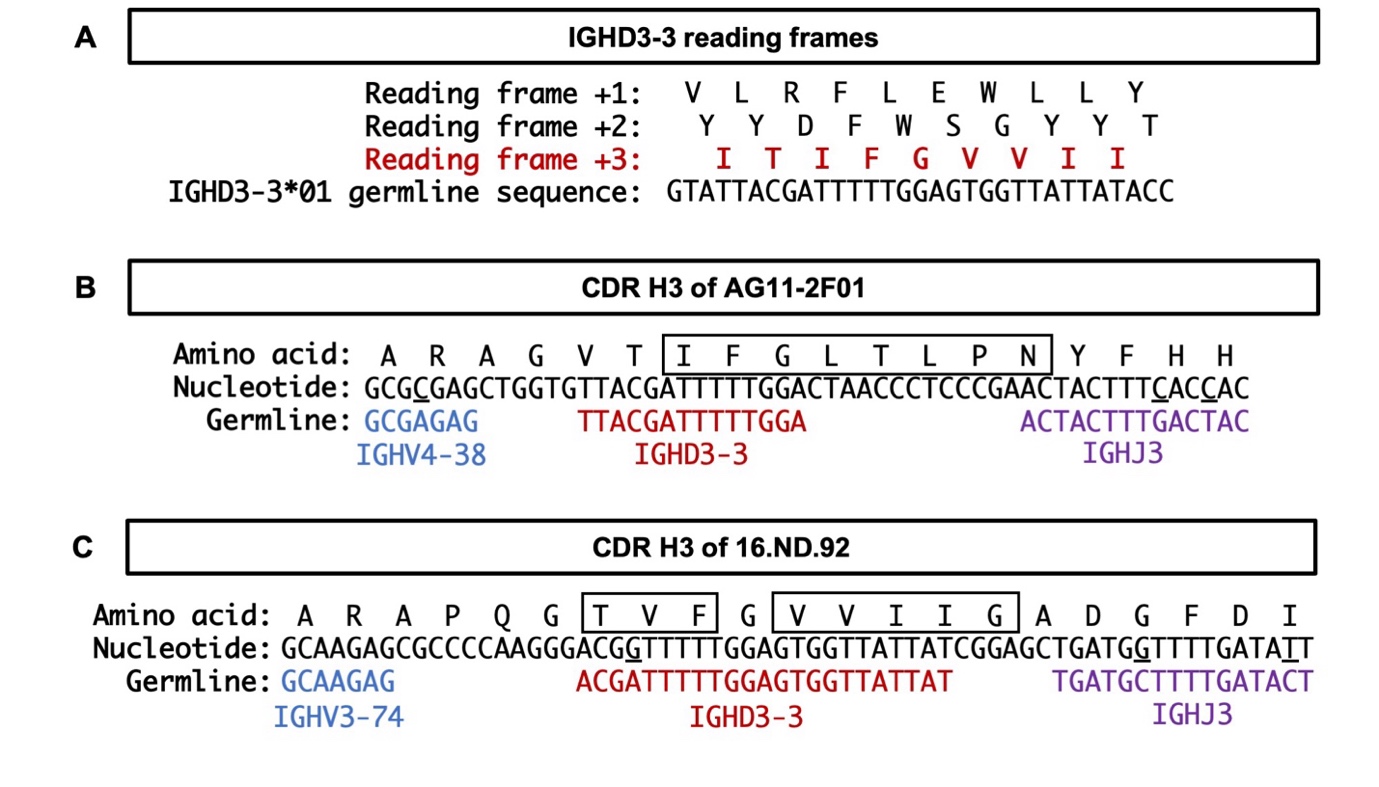

**Figure S13. CDR H3 sequence analysis of AG11-2F01 and 16.ND.92. (A)** The three forward reading frames of the IGHD3-3 are shown with reading frame +3 highlighted in red. **(B-C)** Germline analysis of the CDR H3 sequences of AG11-2F01 and 16.ND.92. Sequences corresponding to the putative IGHV, IGHD, and IGHJ germline genes were annotated by blue, red, and purple, respectively. Somatically mutated nucleotides are underlined. Intervening spaces at the V-D and D-J junctions are N-nucleotide additions. Paratope residues encoded in the CDR H3 are highlighted by the boxes.

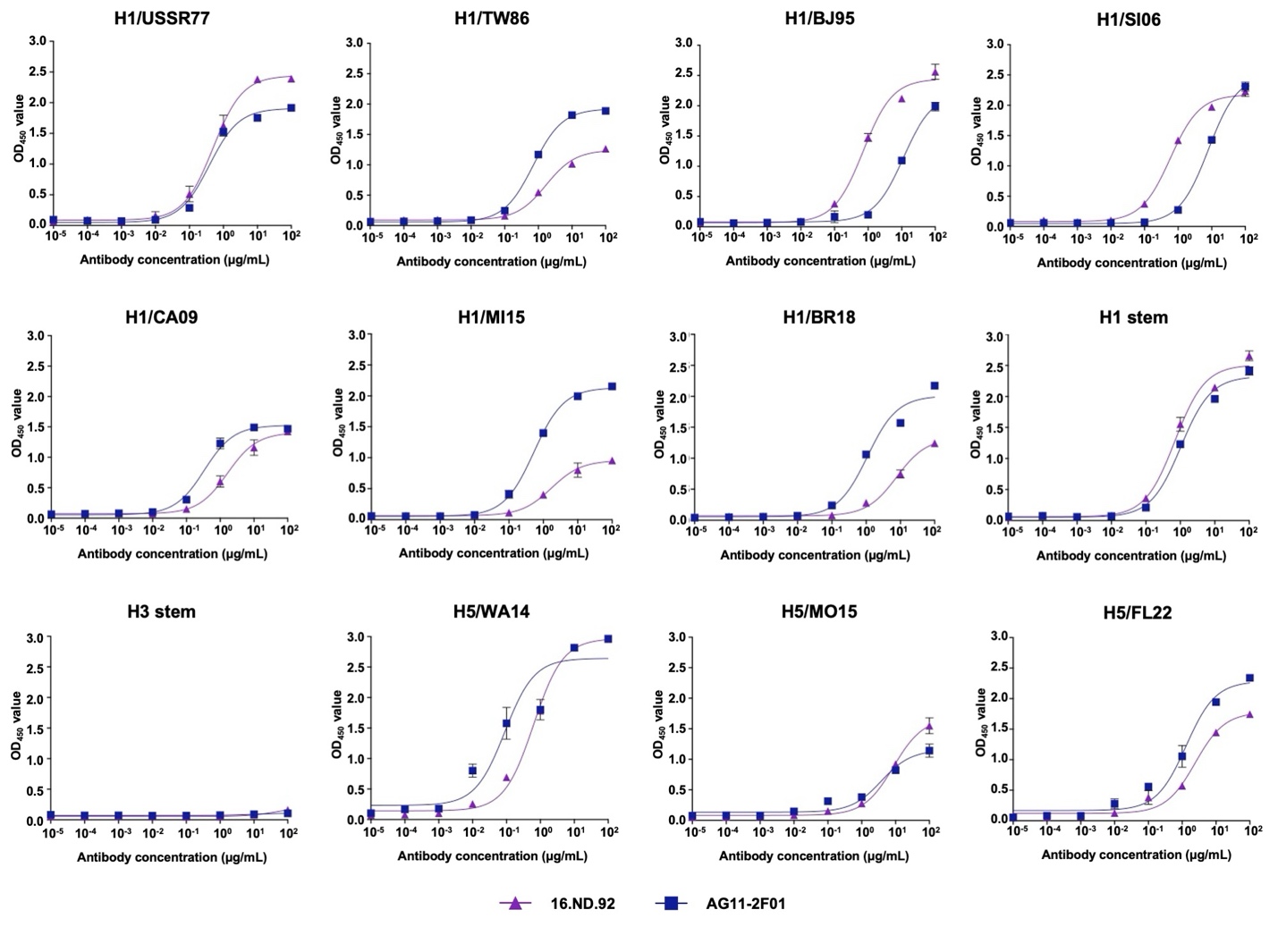

**Figure S14. ELISA titration curves.** The binding activities of AG11-2F01 (blue) and 16.ND.92 (purple) at the indicated concentrations against recombinant HAs were tested by ELISA. The OD_450_ values are plotted as the average of two replicates with the error bars representing the standard deviation.

**Table S5. CR9114 competition data of validated antibodies**

| Antigen | Antibody | Competition Index | BLI competition % |
| --- | --- | --- | --- |
| H1/SI06 | AG11-2F01 | 1.18 | 100 |
| H1/SI06 | 31.a.55 | 0.34 | 99.24 |
| H1/SI06 | 009-10-1G06 | -0.15 | 31.7 |
| H1/SI06 | 045-09-1G05 | -0.39 | 25.7 |
| H1/SI06 | 009-10-2F01 | -0.53 | 30.42 |
| H1/SI06 | 051-10-2B05 | -0.60 | 22.27 |
| H1/MI15 | 31.a.55 | 1.79 | 45.16 |
| H1/MI15 | 029-09-2A06 | -0.22 | 6.25 |
| H1/MI15 | 009-10-1G06 | -0.42 | 0 |
| H1/MI15 | 045-09-1G05 | -0.42 | 0 |
| H1/MI15 | 042-10-2D01 | -0.59 | 8.18 |
| H1/MI15 | 009-10-2F01 | -0.98 | 0 |
| H5/QH05 | 3C11 | 0.22 | 0 |
| H5/QH05 | 56.e.01 | 0.01 | 13.29 |
| H5/QH05 | C7-3-C02 | -0.09 | 0 |
| H7/SH13 | 045-09-1G05 | -1.11 | 0 |
| H7/SH13 | 042-10-2D01 | -1.24 | 0 |
| B/Phu13 | 011-10069-2G04 | 0.03 | 0 |
| B/Phu13 | K77-2A04 | 0.84 | 69.72 |
| B/Phu13 | 017-10116-3D04 | 0.06 | 76.9 |
| B/Phu13 | 011-10069-2G04 | 0.03 | 79.76 |
| B/Phu13 | 034-100809-1F05 | -0.51 | 18.53 |
| B/Lee40 | K77-2A04 | 1.42 | 68.78 |
| B/Lee40 | 011-10069-2G04 | 0.97 | 58.67 |
| B/Lee40 | 017-10116-3D04 | 0.79 | 87.64 |
| B/Lee40 | 150055-015-1D02 | 0.08 | 5.67 |
| B/Lee40 | 034-100809-1F05 | -0.01 | 0 |

**Table S6. Cryo-EM data collection, refinement and validation statistics.**

|  | SI/06 HA + 2F01  (EMDB 46727)  (PDB 9DBX) | SI/06 HA + 16.ND.92  (EMDB 45930)  (PDB 9CU7) |
| --- | --- | --- |
| **Data collection and processing** |  |  |
| Magnification | 130,000 | 81,000 |
| Voltage (kV) | 300 | 300 |
| Electron exposure (e^–^/Å^2^) | 50.00 | 57.35 |
| Defocus range (μm) | -0.8 to -1.5 | -0.5 to -5.0 |
| Pixel size (Å) | 0.66 | 0.53 |
| Symmetry imposed | C3 | C3 |
| Initial particle images (no.) | 1,301,328 | 777,446 |
| Final particle images (no.) | 359,720 | 152,449 |
| Map resolution (Å)  FSC threshold 0.143 | 2.89 | 2.82 |
| Map postprocessing | N/A | DeepEMhancer |
| **Refinement** |  |  |
| Initial model used (PDB code) | 6FYT | 6FYT |
| Model resolution (Å)  FSC threshold |  |  |
| Model resolution range (Å) |  |  |
| Map sharpening *B* factor (Å^2^) | 95.4 | N/A |
| Model composition  Non-hydrogen atoms  Protein residues  Ligands | 17,055  2157 | 16,913  2,172 |
| *B* factors (Å^2^)  Protein  Ligand |  |  |
| R.m.s. deviations  Bond lengths (Å)  Bond angles (°) | 0.003  0.576 | 0.003  0.513 |
| Validation  MolProbity score  Clashscore  Poor rotamers (%) | 1.78  7.87  1.23 | 2.19  6.77  4.76 |
| Ramachandran plot  Favored (%)  Allowed (%)  Disallowed (%) | 95.9  4.1  0.0 | 95.6  4.4  0.0 |

**Table S7: Buried surface areas upon binding of IGHD3-3 antibodies.**

| **Antibody** | **V_H_ BSA (Å^2^)** | | | **V_L_ BSA (Å^2^)** | **Total BSA (Å^2^)** |
| --- | --- | --- | --- | --- | --- |
|  | **CDRH3** | | **Non-CDRH3** |  |  |
|  | **IGHD3-3** | **Other** |  |  |  |
| AG11-2F01 | 379.3 | 90.2 | 103.7 | 349.7 | 922.9 |
| 16.ND.92 | 519.9 | 7.3 | 0 | 439.4 | 966.6 |
| MEDI-8852 | 393.9 | 3.8 | 265.0 | 281.9 | 944.6 |
| 56.a.09 | 357.2 | 0.7 | 206.8 | 374.7 | 939.4 |
| 54-1G05 | 385.3 | 0 | 313.1 | 539.3 | 1237.7 |
| PN-SIA28 | 289.5 | 258.5 | 207.5 | 283.6 | 1039.1 |
| 39.29 | 328.5 | 147.8 | 79.2 | 641.4 | 1196.9 |
| 429 B01 | 254.4 | 262.7 | 104.5 | 288.2 | 909.8 |

**Table S8. Cost and time comparison between traditional approaches for antibody specificity characterization and oPool^+^ display.**

**Traditional approaches:**

| Steps/reagents | Cost per antibody | Cost for 325 antibodies | Time (1 person) |
| --- | --- | --- | --- |
| **Cloning** |  |  |  |
| dsDNA constructs (V_H_ and V_L_) | ~$55-70 | ~$17,875-22,750 |  |
| Plasmid assembly | ~$25-40 | ~$8,125-13,000 | ~2-7 days |
| Transformation | ~$15-25 | ~$4,875-8,125 | ~2-7 days |
| Plasmid extraction  (mini/midi-prep) | ~$10-15 | ~$3,250-4,875 | ~2-7 days |
| Sequencing to confirm correct constructs | ~$20-40 | ~$6,500-13,000 | ~2-5 days |
| **Expression/purification** |  |  |  |
| Expression (cells, transfection reagents, medium) | ~$30-50 | ~$9,750-4,875 | ~7-30 days |
| Purification | ~$10-20 | ~$3,250-6,500 | ~7-30 days |
| **Specificity screen** |  |  |  |
| ELISA (plates, reagents) | ~$85-100 | ~$27,625-32,500 | ~3-10 days |
| **Total** | ~$200-350 | ~$70,000-110,000 | Weeks to months |

**oPool^+^ display:**

| Steps/reagents | Cost per antibody | Cost for 325 antibodies | Time (1 person) |
| --- | --- | --- | --- |
| **Library assembly** |  |  |  |
| Oligo pools | $25.33 | $8,233.48 |  |
| PCR assembly (KAPA HiFi) | $0.13 | $43.31 | 6-10 hours |
| **mRNA display** |  |  |  |
| *In vitro* transcription | $0.12 | $38.18 | 4-16 hours |
| Puromycin ligation | $0.15 | $47.95 | 2 hours |
| *in vitro* translation | $0.16 | $53.10 | 2-3 hours |
| FLAG purification | $0.08 | $24.72 | 2-3 hours |
| Reverse transcription | $0.04 | $12.17 | 0.5 - 1 hour |
| Selection | $0.46 | $150.79 | 1.5- 3 hours |
| **Sequencing preparation** |  |  |  |
| Recovery & barcode PCR (PrimeSTAR Max) | $0.11 | $35.70 | 3 hours |
| **Next Generation Sequencing** |  |  |  |
| Revio SMRTcell 8M | $7.41 | $2,410 |  |
| **Total** | $33.99 | $11,049.4 | 3-5 days |

**Table S9. Sequences of primers used in this study.**

| **Name** | **Sequences** |
| --- | --- |
| oPool_assembly_F | 5’-TTC TAA TAC GAC TCA CTA TAG GGA CAA TTA CTA AAG GAG TAT CC-3’ |
| oPool_assembly_R | 5’-GGA GCC GCT ACC CTT ATC GTC GTC ATC CTT GTA ATC GGA TCC T-3’ |
| Splint_oligo | 5’-TTT TTT TTT TTT GGA GCC GCT ACC-3’ |
| Puromycin_linker | 5’-/5Phos/-d(A)21-(C_9_)3-d(ACC)-puromycin-3’ |
| Oligo(dT)_21_ | 5’-TTT TTT TTT TTT TTT TTT TTT -3’ |
| oPool_recovery_F | 5’-GTA AAA CGA CGG CCA GTT TCA GGG GAC AAT TAC TAA AGG AGT ATC C-3’ |
| oPool_recovery_R | 5’- CAG GAA ACA GCT ATG ACC CAC TCG TCA TCC TTG TAA TCG GAT CCT CCG GA-3’ |
| PacBio_F1 | 5’-TGA ACC TTG TAA AAC GAC GGC CAG TTT CAG-3’ |
| PacBio_F2 | 5’-TGC TAA GTG TAA AAC GAC GGC CAG TTT CAG -3’ |
| PacBio_F3 | 5’-TGT TCT CTG TAA AAC GAC GGC CAG TTT CAG -3’ |
| PacBio_F4 | 5’-TAA GAC ACG TAA AAC GAC GGC CAG TTT CAG-3’ |
| PacBio_F5 | 5’-CTA ATC GAG TAA AAC GAC GGC CAG TTT CAG-3’ |
| PacBio_F6 | 5’-CTA GAA CAG TAA AAC GAC GGC CAG TTT CAG-3’ |
| PacBio_R8 | 5’-GTG TGG TGC AGG AAA CAG CTA TGA CCC ACT -3’ |
| PacBio_R9 | 5’-TGG GTT TCC AGG AAA CAG CTA TGA CCC ACT -3’ |

**Table S10. Custom cutoff used for the analysis of each screen.**

| Screen | Binding score cutoff |
| --- | --- |
| H1 stem | 28 |
| H3 stem | 28 |
| H1/SI06 | 5 |
| H1/SI06+CR9114 | 9 |
| MI15/H1 | 3 |
| MI15/H1+CR9114 | 7 |
| SP16/H3 | 5 |
| H5/QH05 | 19 |
| H5/QH05+CR9114 | 10 |
| H7/SH13 | 2.85 |
| H7/SH13+CR9114 | 10 |
| B/Phu13 | 5 |
| B/Phu13+CR9114 | 10 |
| B/Lee40 | 5 |
| B/Lee40+CR9114 | 11.5 |

| Competition Index | Index cutoff  over: CR9114 competing  under: CR9114 non-competing |
| --- | --- |
| H1/SI06 | log_10_(2) |
| H1/MI15 | log_10_(2.5) |
| H5/QH05 | log_10_(2.5) |
| H7/SH13 | log_10_(2) |
| B/Phu13 | log_10_(1.65) |
| B/Lee40 | log_10_(1.4) |
